# Neuronal Maturation-dependent Nano-Neuro Interaction and Modulation

**DOI:** 10.1101/2022.08.03.502650

**Authors:** Prashant Gupta, Priya Rathi, Rohit Gupta, Harsh Baldi, Quentin Coquerel, Avishek Debnath, Hamed Gholami Derami, Baranidharan Raman, Srikanth Singamaneni

## Abstract

Nanotechnology-enabled neuromodulation, a rapidly growing technique, is a promising minimally-invasive tool in neuroscience and engineering for both fundamental studies as well as clinical applications. However, the nano-neuro interactions at different stages of maturation of a neural network and its implications on the nano-neuromodulation remain unclear. Here, we report heterogeneous to homogenous transformation of neuromodulation in a progressively maturing neural network. Utilizing plasmonic fluors as ultrabright fluorescent nanolabels, we reveal that negative surface charge of the nanoparticles renders selective nano-neuro interaction with a strong correlation between the maturation stage of the individual neurons in the neural network and the density of the nanoparticles bound on the neurons. In stark contrast to homogeneous neuromodulation in a mature neural network reported so far, the maturation-dependent density of the nanoparticles bound to neurons in a developing neural network resulted in a heterogeneous optical neuromodulation (i.e., simultaneous excitation and inhibition of neural network activity). This study advances our understanding of nano-neuro interactions and nano-neuromodulation with potential applications in minimally-invasive technologies for treating neuronal disorders in parts of mammalian brain where neurogenesis persists throughout aging.

---

One of the major goals of modern biomedical research is to understand the working principles of nervous system.^1^ As a rapidly growing technique, neuromodulation has proved to be of paramount importance in answering fundamental neuroscience questions and in devising advanced treatments of various neurological disorders.^2^ Electrical neuromodulation-based implantable devices developed over the past few decades have proved effective in the treatment of many debilitating medical conditions including Parkinson’s disease, clinical depression, and epilepsy.^3^ However, the use of these devices (usually metal electrodes), owing to their bulkiness, mechanical invasiveness and inability to target individual neurons and neuronal circuits, are often limited.^4, 5^ Optogenetics, involving genetic modification to control cellular activity via optical stimuli, has emerged as an attractive alternative tool over the past two decades.^6, 7^ Although, optogenetics overcomes many of the aforementioned issues associated with physical electrodes,^7, 8^ it relies on genetic modification of neurons, which is irreversible, difficult to implement in model organisms without rich repertoire of genetic tools and difficult to translate to humans.^9^ As such, nanomaterials based non-genetic neuromodulation approaches, which can be administered in a drug-like fashion, have been explored in recent years.^1, 10^ Nano-enabled neuromodulation involves harvesting energy from an external source by the nanomaterials in a wireless manner and transducing it into physiologically-relevant stimuli in a localized region (down to single neurons or subcellular compartments) for neural stimulation.^11^ Nano-neuromodulation also provides additional flexibility towards stimulation modes based on the energy sources employed in conjunction with specific transducing nanostructures such as optical^12^, acoustic^13^ and magnetic^14^ stimulation. Among these, optical stimulation via photothermal nano-transducers (such as plasmonic nanostructures, graphene, polydopamine nanoparticles, etc.) have shown great promise and versatility.^15-23^

Majority of the photothermal neuromodulation studies involve primary neuron culture close to its complete maturation stage as the model system. Although, in most brain regions, neurogenesis (process of generating new functional neurons from precursors) has been confined to a discrete developmental period, life-long neurogenesis has been observed in both the hippocampus and subventricular zone of almost all mammals, including humans.^24^ The addition of new neurons to the complex circuitry of adult brain plays crucial role in memory and behavior.^25^ Interestingly, these immature neurons exhibit high excitability, reduced GABAergic inhibition and a lower threshold for the induction of long-term potentiation, which allows them to spike despite their developing glutamatergic inputs and participate in information processing before reaching a complete maturation stage.^25^ The interaction of nanomaterials with these young neurons, if any, in the heterogeneous neural network comprising of both young and mature neurons, and its implications on the nano-neuromodulation is yet to be elucidated. This improved understanding would pave the way in designing minimally-invasive non-genetic nanomaterial-based neuromodulation techniques for both fundamental studies and clinical applications.

Recently, Dante et al. described the critical role of the surface charge of nanoparticles in their selective binding to neurons.^26^ They demonstrated that negatively charged nanoparticles, irrespective of shape, size and material composition of the nanomaterial, exclusively bind to excitable neuronal cells and never to non-excitable glial cells whereas positively charged and neutral particles never spontaneously bind to neurons. In this study, we harness plasmonic fluors-IR650 (PFs), ultrabright fluorescent nanoconstructs recently developed by our research group,^27^ to unveil the neuronal maturation-dependent nano-neuro interactions (**Figure 1**). Building on these findings, we rationalize the nongenetic optical neuromodulation in both heterogeneous neural network (comprising of both young and mature neurons) and homogeneous neural network (majorly comprising of mature neurons) utilizing a commonly employed plasmonic photothermal nanotransducer, gold nanorods.^15, 17, 19, 21, 22^

**Figure 1.**
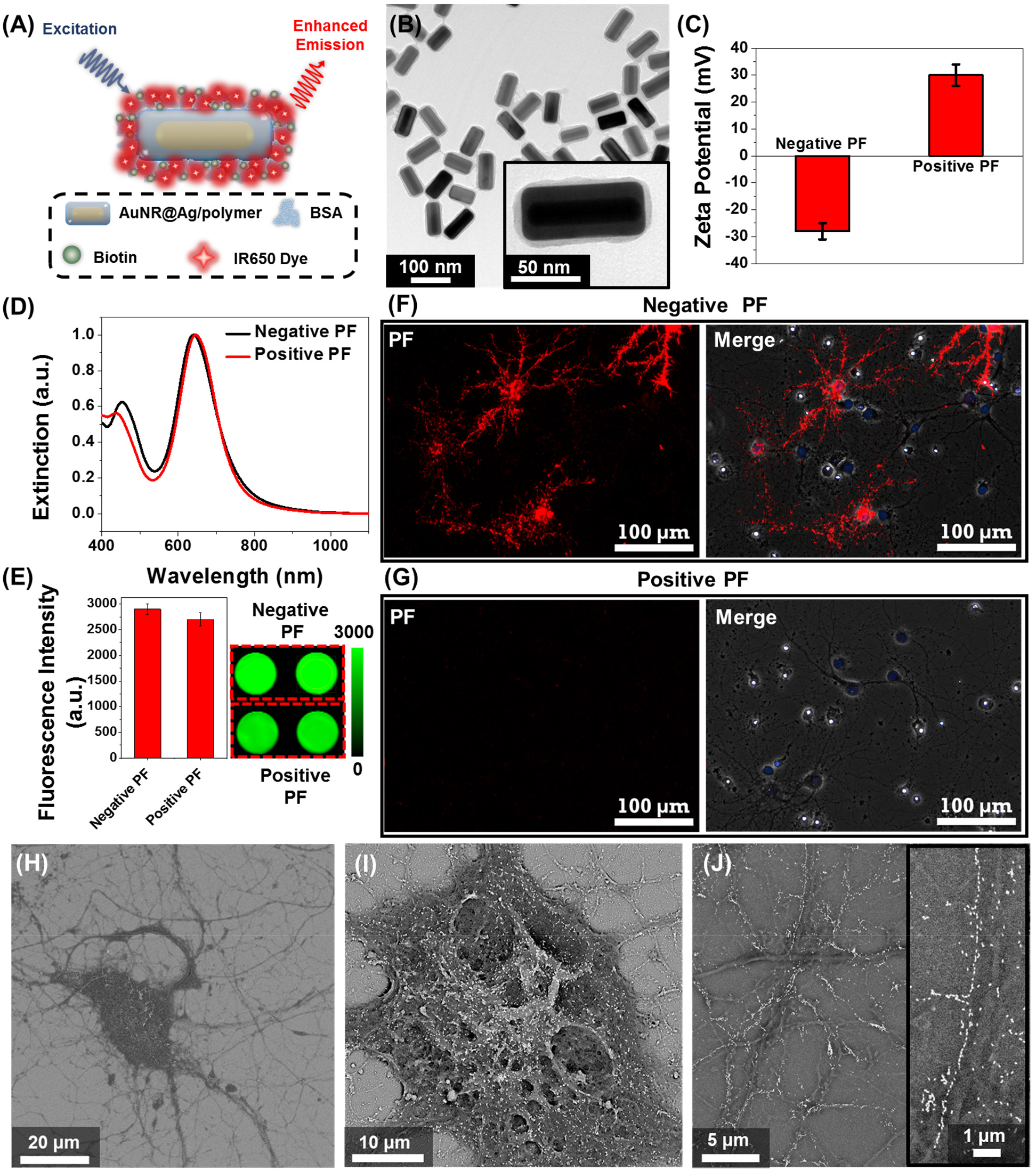
Plasmonic-fluor as an ultrabright fluorescent nanoconstruct for understanding nano-neuro interaction. (A) Schematic illustration of plasmonic-fluor (PF) comprised of plasmonic nanoantenna (Au@Ag nanocuboid) coated with a polymer layer as dielectric spacer (polymer), fluorophores (IR-650) and a universal biorecognition element (biotin) assembled using bovine serum albumin (BSA). (B) TEM image of PFs (Inset: Higher magnification image depicting a thin organic layer around the plasmonic core). (C) Zeta potential (Error bars, s.d., n = 3 repeated tests), (D) visible–NIR extinction spectra, and (E) Fluorescence intensity (Error bars, s.d., n = 4 independent tests) of negatively and positively-charged PF. Confocal fluorescence images of cultured hippocampal neurons at DIV 14 after 1 hour incubation with (F) negative and (G) positive PFs (red). The nucleus was stained with DAPI (Blue) post-fixation. This is a representative image from 1 of a total of 8 images taken from n=2 independent experiments. SEM image of (H) a single hippocampal neuron with selective localization of negative PFs and a higher magnification image showing (I) the randomly oriented PFs on soma and (J) the longitudinally aligned PFs on the neurites (Inset: zoomed in image depicting single nanoparticle-wide PFs along the neurites). This is a representative image from 15 images taken from n=2 independent experiments.

## Results

### Role of the surface charge of nanoparticles in binding to neurons

We employed plasmonic-fluors comprised of a near infrared dye IR-650 (PF-650) as model nanostructures to understand the interactions between nanoparticles and neurons. We have recently introduced plasmonic-fluors as ultrabright fluorescent nanoconstructs that are nearly 7000-fold brighter compared to the corresponding molecular fluorophores.^27^ PF-650 is comprised of Au@Ag nanocuboids as plasmonic nanoantenna, siloxane copolymer layer as dielectric spacer and BSA-biotin-IR650 conjugates (**Figure 1A**). Transmission electron microscopy (TEM) image depicts the Au@Ag nanocuboids with a length 98 ± 5 nm and a width 42 ± 2.5 nm and the polymer and BSA-biotin-IR650 coating of 3 ± 1 nm (**Figure 1B**). Owing to the presence of the BSA on the surface, under physiological pH conditions, the PFs are negatively charged (with ζ-potential of - 28 ± 3 mV), henceforth termed as negatively-charged PFs (**Figure 1C**). The positively charged PFs (with ζ-potential of +30 ± 4 mV) were obtained by coating negative PFs with poly(allylamine hydrochloride). The positively charged PFs showed no sign of aggregation as evidenced by the absence of broadening of the localized surface plasmon resonance (LSPR) band in the extinction spectrum and the retained florescence intensity (**Figure 1D, E**). To investigate the interaction of nanoparticles with the neurons in an electrically-active neuronal network, primary hippocampal neuronal culture at DIV 14 was incubated with the negative and positive PFs for 1 hour in NbActiv 4 medium, which is a serum-free medium. The absence of serum in the medium precludes the formation of protein corona on the nanoparticles, thus preserving their surface state. We observed that the negatively charged PFs readily bind to the neurons as evidenced by the co-localization of the PF fluorescence signal (λ_emission_ = 650 nm) with the neurons (**Figure 1F**). On the other hand, the positively charged particles do not bind to the neurons (**Figure 1G**), as reported previously.^26^ This observation suggests that the negative surface charge is a necessary condition for the spontaneous binding (*i*.*e*., without any specific targeting moiety) of the nanoparticles to the neurons. The PFs uniformly decorated the soma and the neurites of the neurons. The anisotropic nanostructures (*i*.*e*., nanocuboids) bound on the soma and thicker regions of neurite exhibited random orientation whereas those bound on the thinner regions of neurites were oriented along the length of the neurites (**Figure 1H, I, J**). Notably, in most cases, the PFs bound to the thinner region of neurites formed a single-particle wide linear array. Considering that the lateral dimensions of the neurites is 100 - 1000 nm, the longitudinal alignment of PFs possibly stems from the maximal interfacial contact area of the PFs with the neurites under this orientation.^28-30^

### Effect of nanoparticle binding on the electrical activity of neural network

While PFs are extremely bright fluorescent nanoconstructs that serve as ideal nanolabels to monitor the binding of the nanostructures to neurons, they are not commonly employed for optical neuromodulation. Owing to the facile tunability of the LSPR wavelength over a broad range and their large extinction cross-section, gold nanorods (AuNRs) are highly attractive photothermal nanotransducers for optical neuromodulation.^15, 17, 19, 21, 22^ We set out to investigate the effect of the binding of the negatively-charged AuNRs on the electrical activity of the neurons (Figure S1A). Negatively charged AuNRs were obtained by coating the as synthesized AuNRs with polystyrene sulfonate (PSS). Following the PSS coating, the AuNRs exhibited a blue shift (of 10 nm) in the LSPR wavelength and a zeta-potential of -34 ± 4 mV (Figure S1B). To investigate the change in the electrical activity of neurons in response to the nanoparticle binding, hippocampal neurons were cultured on microelectrode arrays (MEAs) consisting of 60 electrodes. Extracellular activity of neurons was recorded with and without AuNR incubation (**Figure 2A**). Neurons cultured on MEA formed a dense network of neurites around TiN recording electrodes (Figure S2). To investigate the effect of nano-neuro interaction on the electrical activity of the primary hippocampal cultured neurons, the extracellular activity was recorded for 10 min prior to AuNR binding at 14 *days-in-vitro* (DIV 14) (**Figure 2B**). The neurons cultured on the MEAs were then incubated with AuNRs (at a final concentration of optical density (O.D.) ∼ 0.5) for 1 hour followed by rinsing with medium. Following the binding of AuNRs to the neurons, although there is a significant change in the spontaneous electrical activity of the neuronal network, the spike shape and amplitude remained unaltered (**Figure 2B, C, S3 and S4**). The mean spike rate of the network measured over a period of 10 min reduced significantly while the burst activity significantly increased (**Figure 2D, E**). Note that the duration of the burst events and number of spikes per burst decreased significantly after binding of the AuNRs (**Figure 2F, G**).

**Figure 2.**
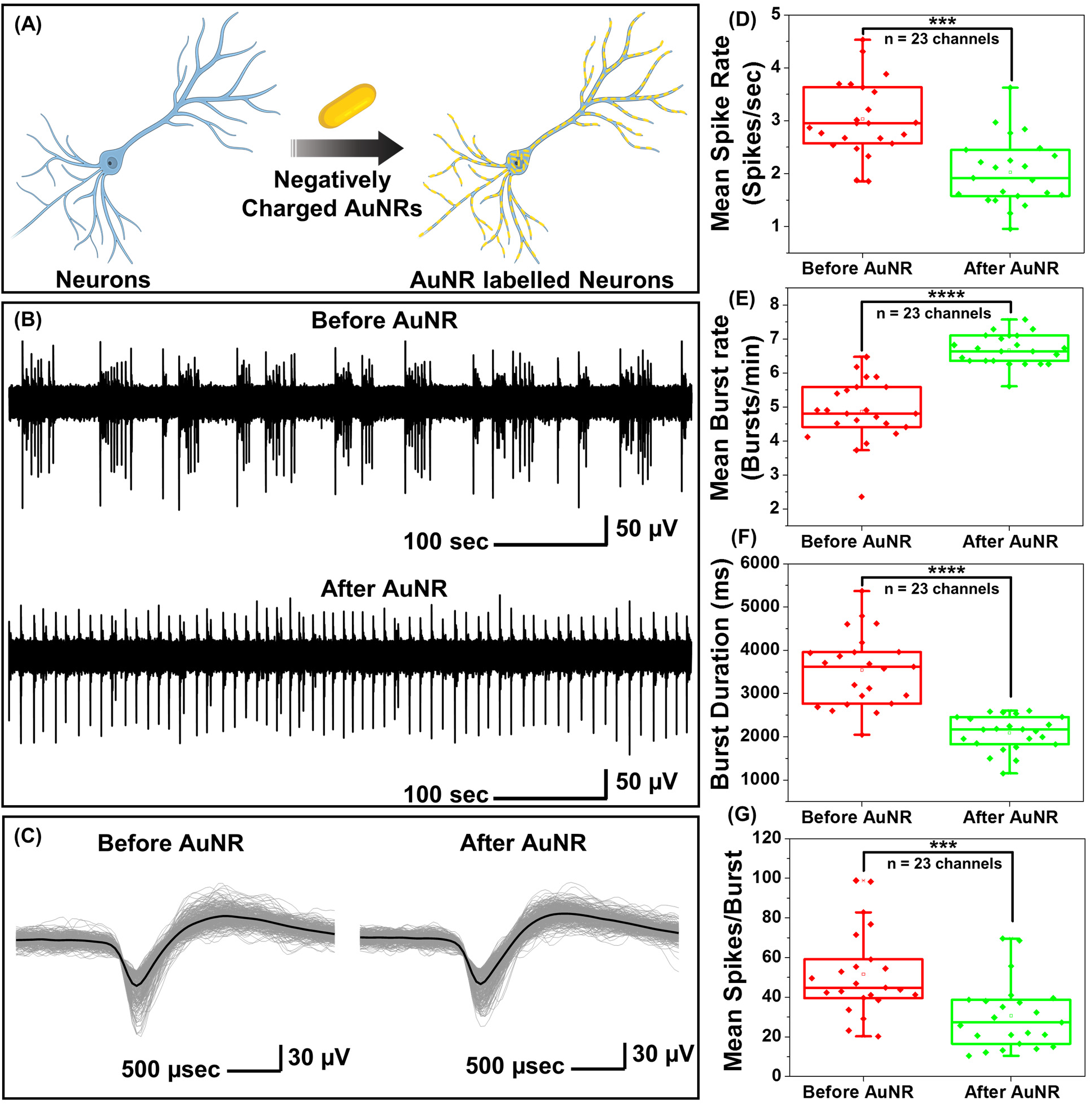
Nano-neuro interaction elicits electrophysiological alterations in *in-vitro* cultured hippocampal neurons. (A) Schematic illustration depicting the selective binding of negatively charged plasmonic nanostructures (gold nanorods, AuNR) to hippocampal neurons. (B) A single trace of spike recording before and after neurons were incubated with negatively charged AuNR. (C) Overlaid spike waveform of hippocampal neurons before and after AuNR labeling. Panel on the left shows the spike cutouts before the application of AuNRs and panel on the right shows the spike cutouts after the AuNR binding. Spikes from 10-minute recording with at least 700 spikes in each set. Black curve shows the mean value for each set. The traces in B and C are representative ones from a total of 23 active channels measured from primary cultured hippocampal neurons cultured on a microelectrode array (MEA). The experiment was repeated three times independently with similar results. Effect of AuNR localization on neuron membrane on the (D) mean spike rate (Unpaired Two-samples t-test; p= 0.0002, n=23, * p<0.05, ** p<0.01, *** p<0.001 and **** p<0.0001), (E) mean burst rate (Unpaired Two-samples t-test; p< 0.0001, n=23, * p<0.05, ** p<0.01, *** p<0.001 and **** p<0.0001), (F) burst duration (Unpaired Two-samples t-test; p< 0.0001, n=23, * p<0.05, ** p<0.01, *** p<0.001 and **** p<0.0001) and (G) mean spikes per burst (Unpaired Two-samples t-test; p= 0.0006, n=23, * p<0.05, ** p<0.01, *** p<0.001 and **** p<0.0001) of cultured neurons.

### Homogenous and heterogeneous modulation of neuronal activity through photothermal stimulation

Under near infrared laser illumination, the plasmonic nanostructures bound on the neurons result in localized temperature rise, which in turn either reversibly alters the electrical capacitance and therefore the excitability of the neurons or reversibly activates the temperature sensitive TREK-1 ion channels and consequently reduces the discharge of action potentials.^17, 31, 32^ Owing to their ability to readily bind to neurons (**Figure 1F**), we employed negatively charged AuNRs to understand the effect of maturation stage of neuronal network on the photothermal neuromodulation. Hippocampal neurons cultured on MEAs were incubated with negatively charged AuNRs (76.2 pM final concentration) for 1 hour at DIV 14, 18, 22 and 26, followed by washing with the NbActiv4 medium. Different MEAs were utilized at different DIVs, so as to avoid any interference from nanoparticle-induced neuronal membrane depolarization on the neuron maturation process.^26^ Note that the kinetics of the neuronal maturation is significantly modulated by local network activity.^33^

The AuNR localized neurons were subjected to repeated irradiation of 808 nm laser at laser power density of 14 mW/mm^2^ for different durations (10, 20, 30 and 60 seconds) in a back-to-back pulsatile fashion (**Figure 3A**). The extracellular activity of the neurons was recorded before, during, and after the photothermal treatment. The extracellular signal recorded by each of MEA channels corresponds to a group of neurons on and around the channels that are irradiated by the NIR laser. At DIV 14 and 18, a small fraction of channels (10 to 30%) exhibited complete inhibition of neural activity in response to photothermal stimulation (observed from mean spike rate change before, during and after laser illumination) which may be attributed to the membrane-localized photothermal heating via AuNRs. However, most of the electrodes depicted partial reduction, enhancement or no change in spiking activity (**Figure 3B, D**). In the channels where spiking activity was suppressed during photothermal treatment, the shape and amplitude of the remnant spikes remained unaltered before and after laser illumination suggesting the reversibility of the neuromodulation (**Figure 3C**, top panel). In addition, no significant difference in the spike shape and amplitude was observed in the channels exhibiting excitation during laser stimuli (**Figure 3C**, bottom panel). Furthermore, for all cells, no significant difference in the mean spike rate was observed before and after photothermal neuromodulation, which further confirms the complete reversibility of the nano-neuromodulation (Figure S5). At DIV 22 and 26, the photothermal neuromodulation resulted in nearly complete inhibition of spiking activity (**Figure 3B, D**). Moreover, with an increase in DIV from 14 to 18, a larger fraction of the electrodes exhibited inhibition in response to photothermal stimulation, finally reaching 100% at DIV 22 and above (**Figure 3E**). These results highlight the transformation of the photothermal neuromodulation from a heterogeneous (in early stages of DIV 14 and 18) to homogeneous (in later stages of DIV 22 and 26) change in electrical activity.

**Figure 3.**
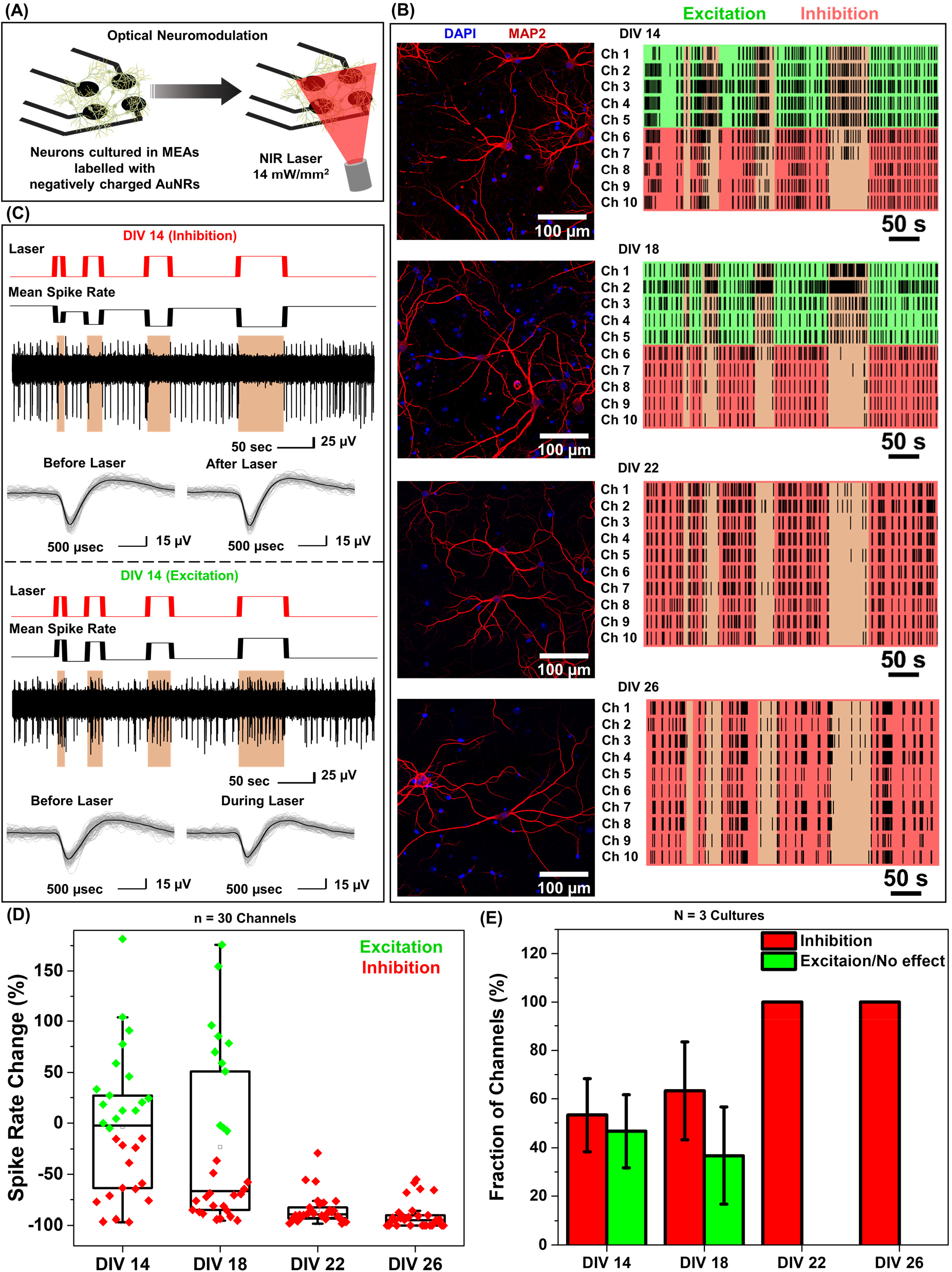
Homogenous and heterogeneous modulation of neuronal activity through photothermal stimulation. (A) Schematic illustration of the optical neuromodulation experimental setup demonstrating primary hippocampal neurons cultured in MEAs and stimulated with NIR laser (808 nm, 14 mW/mm^2^) after incubation with negatively charged AuNRs. (B) Raster plots (right panel) representing the spiking activity of primary hippocampal neurons labeled with negatively charged AuNRs at different days in vitro (DIV 14, 18, 22 and 26). Each row in the raster plot corresponds to one channel of a MEA. Ten representative channels out of at least 30 active channels are presented. The vertical orange color bar indicates the time when NIR laser (808 nm laser wavelength, 14 mW/mm^2^ power density, laser duration of 10, 20, 30 and 60 seconds) was illuminated on the MEAs with primary hippocampal neurons labelled with negatively charged AuNRs to investigate optical neuromodulation. The experiment was repeated three times independently with similar results. Corresponding confocal fluorescence images (left panel) of primary cultured hippocampal neurons at DIV 14, 18, 22 and 26 co-stained with MAP2 (red) which is a neuronal marker and DAPI (blue) for nucleus staining. (C) Raw extracellular voltage traces showing modulation of spiking activity (top panel in each block) recorded from two different channels, one exhibiting inhibition (top panel) and the other showing excitation (bottom panel) of neural activity in response to optical stimuli measured simultaneously from the MEA with cultured hippocampal neurons at DIV 14. Overlaid spike waveform (bottom panel in each block) of hippocampal neurons before (inhibition and excitation panel), after (inhibition panel) and during (excitation panel) optical neuromodulation (the spikes waveforms are plotted for before, during and after 60 second laser illumination, with at least 90 spikes in each set and black curve shows the mean value for each set). The traces are representative ones from a total of at least 30 active channels measured from primary cultured hippocampal neurons cultured on a MEA. (D) Whisker plot demonstrating the quantification of spike rate changes in panel B (effect of neuronal network maturation on the optical neuromodulation, transformation from heterogeneous to homogeneous neuromodulation, n ≥ 30 channels). (E) Fraction of MEA channels exhibiting inhibition and excitation/no change in the spike rate of the neurons labeled with negatively charged AuNRs in response to NIR stimuli (Error bars, s.d., N = 3 independent cultures).

### Heterogeneous nano-neuro interaction

We hypothesized that the heterogeneity in the nano-neuromodulation at early stages of the neuronal network maturation is associated with the heterogeneity in the nano-neuro interaction. To test this hypothesis, we examined the binding of negative PFs to neurons at DIV 14, a time point at which we observed heterogeneous neuromodulation (**Figure 3E**). We observed that only a small fraction of the cells are tagged with negative PFs while most of the cells are devoid of nanoparticles (**Figure 4A**, untagged cells indicated by yellow arrows). This observation suggested that at DIV 14, there is indeed a heterogeneous binding of nanoparticles in the neural network. We investigated the viability of the cells that are not tagged with negative PFs at DIV 14 using Calcein AM and ethidium homodimer staining (*i*.*e*. live/dead cells assay). We found that even the cells that are not tagged with negative PFs are alive, as confirmed by the confocal fluorescence images (**Figure 4C**, yellow arrows indicate untagged viable cells). We then employed MAP2 as neuronal marker^34^ and Nestin as progenitor cell marker of both neuronal and glial lineage^35^ to differentiate between neurons and glial cells in the culture (neuronal cells expressed both MAP2 (red) and nestin (green) markers while glial cells expressed only nestin (green) marker) and subsequently investigated the presence of unlabeled neurons in the cultured neural network at DIV 14. We observed that a significant fraction of untagged cells (absence of PFs, cyan) expressed MAP2 (red), confirming the heterogeneous binding of negatively charged nanoparticles to the cultured neural network at earlier stages (DIV 14) (**Figure 4D**).

**Figure 4.**
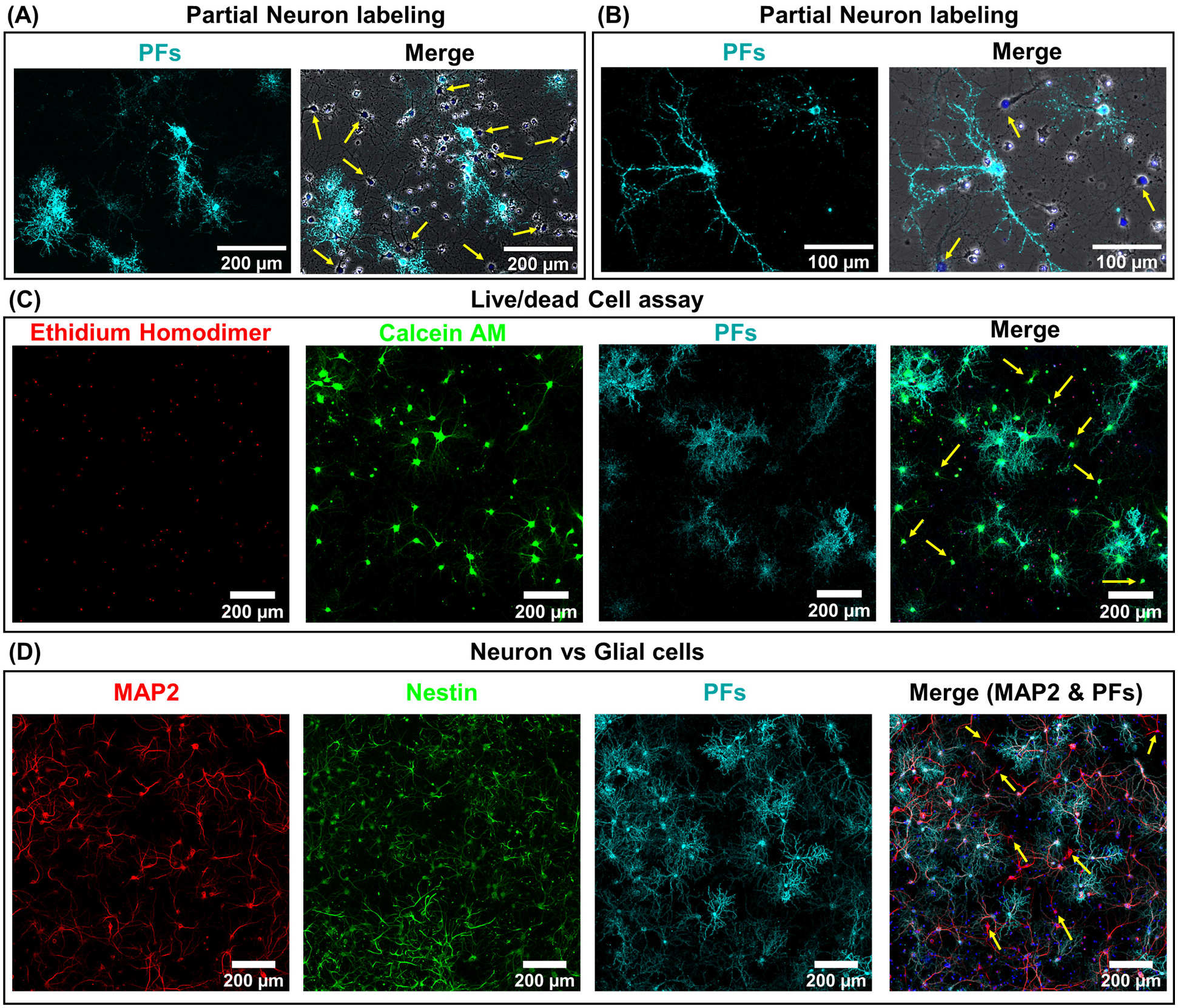
Partial labeling of neurons with negatively charged PFs. (A) Low and (B) high magnification confocal fluorescence images of cultured hippocampal neurons after 1 hour incubation with negatively charged PFs at DIV 14. The left panel shows the fluorescence image corresponding to negatively charged PFs (cyan) and right panel is the merged fluorescence image comprising of phase contrast (gray), DAPI for nucleus staining (blue) and PFs (cyan). Yellow arrows indicate untagged cells. (n=2 independent experiments) (C) Confocal fluorescence images of cultured hippocampal neurons after 1 hour incubation with negatively charged PFs (cyan) at DIV 14, co-stained with ethidium homodimer (red) for dead cell staining and calcein AM (green) for live cell staining. The yellow arrows indicate live cells that are not tagged with negatively charged PFs. (n=2 independent experiments) (D) Confocal fluorescence images of cultured hippocampal neurons after 1 hour incubation with negatively charged PFs (cyan) at DIV 14, co-stained with MAP2 (red, neuronal cell marker, specific to neuron cells), Nestin (green, progenitor cell marker, stains both neurons and glial cells) and DAPI (blue, nucleus staining). The yellow arrows indicate untagged neurons after incubation with negatively charged PFs. (n=2 independent experiments)

### Neuron maturation-dependent nano-neuro interaction

Based on the observation that a significant fraction of the live neurons remain untagged at DIV 14, we hypothesized that this heterogeneity in the nano-neuro interaction is responsible for the heterogeneous neuromodulation in earlier stages (DIV 14 and 18, Figure 3) and homogeneous neuromodulation in later stages (DIV 22 and 26, Figure 3). To test this hypothesis, we employed negative PFs to investigate the neuronal maturation-dependent nano-neuro interaction. We monitored the binding of the negatively charged PFs to neurons at various DIVs (DIV 3, 5, 7, 10, 14, 18, 22 and 26) (**Figure 5A**, individual channels of fluorescence images presented in **Figure S7-S22**). After nanoparticle binding, the neurons were co-stained with MAP2 and nestin post-fixation to distinguish neuronal cells from glial cells. We did not observe discernable binding of negatively charged nanostructures to neurons till DIV 5, suggesting that the nanostructures do not interact with young neurons. However, as the DIV increases above 7, the fraction of neurons tagged with the negatively charged PFs (cyan) and the florescence intensity (representing the number of PFs bound to the neurons) associated with the tagged neurons increased (**Figure 5B, C**). The progressive increase in the nanoparticle binding to the neurons with an increase in DIV may be attributed to the progressive transformation of the neuronal network from young to developing to a finally mature state. Considering that the neuron maturation process is heterogeneous in nature,^36^ young, developing and mature neurons co-exist over the DIV range studied here. However, with an increase in DIV the fraction of young neurons decreases and that of developing neurons and mature neurons increases, until all the neurons in the network mature. These observations suggest that neuron maturation plays a critical role in nanoparticle binding in the *in vitro* neural network, which in turn affects the nano-neuromodulation. We believe that the transformation from heterogeneous response to photothermal stimulation at early stages of the network (DIV 14 and 18) to homogeneous response at later stages (DIV 22 and 26) is a direct manifestation of maturation-dependent tagging of neurons with negatively charged plasmonic nanostructures (**Figure 3**).

**Figure 5.**
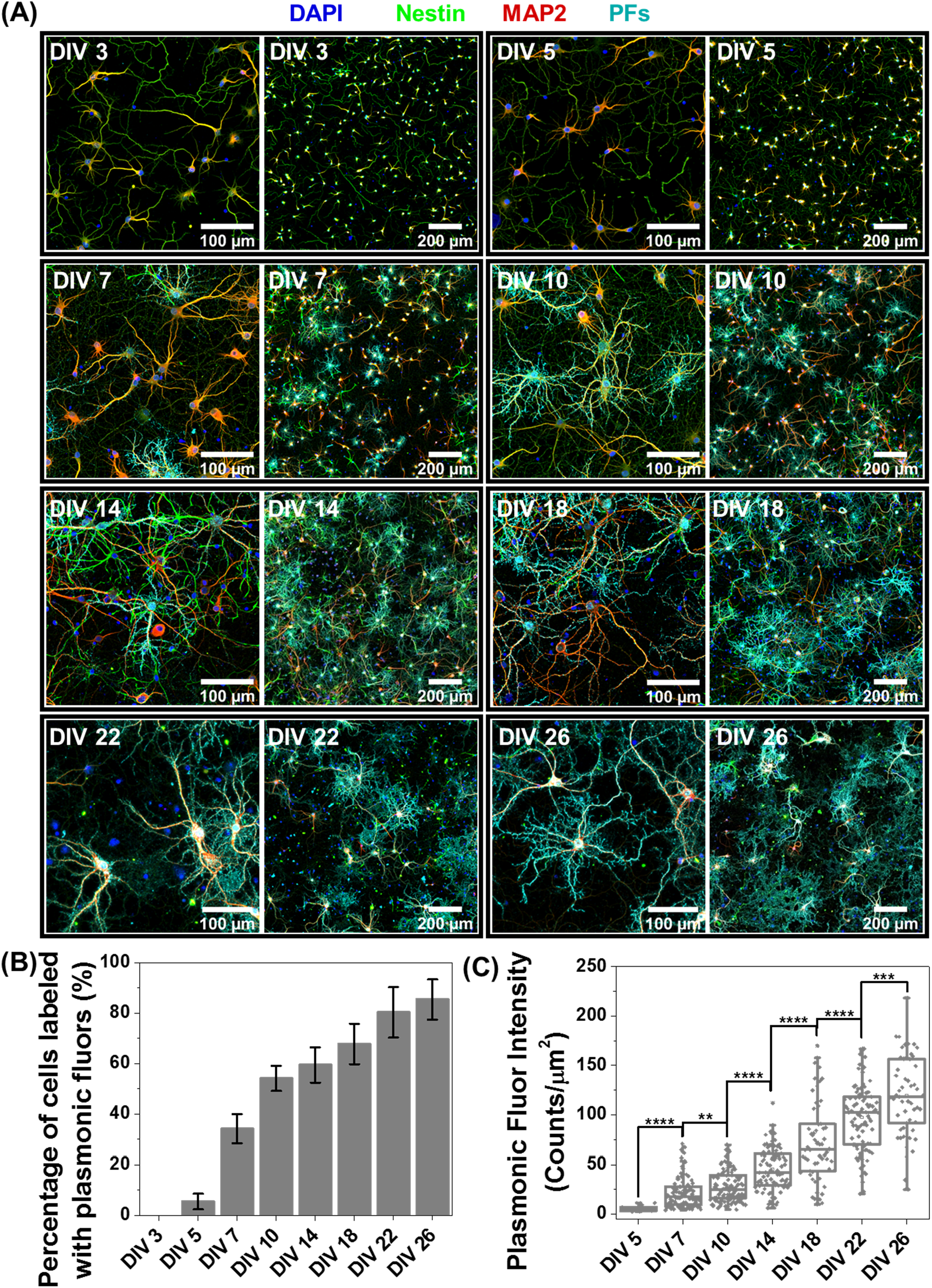
Role of Neuronal network maturation in nano-neuro interaction. (A) Confocal fluorescence images of cultured hippocampal neurons after 1 hour incubation with negatively charged PFs (cyan) at DIV 3, 5, 7, 10, 14, 18, 22 and 26, co-stained with MAP2 (red, neuronal cell marker, specific to neuron cells), Nestin (green, progenitor cell marker, stains both neurons and glial cells) and DAPI (blue, nucleus staining). The left panel in each block shows the fluorescence image at 20X magnification and the left panel each block shows the 3×3 tiled image obtained from 9 images similar to the left panel. (n=2 independent experiments). (B) Percentage of neuronal cells labelled with negatively charged PFs at different DIVs (Error bars, s.d., n = 6, 3×3 tiled images from n=2 independent cultures). (C) Whisker plot representing fluorescence intensity of PF tagged neurons at various DIVs (Unpaired Two-samples t-test; n = 5, 106, 111, 94, 98, 95 and 54 labelled neuronal cells from three 3×3 tiled images from the same culture for DIVs 5, 7, 10, 14, 18, 22 and 26 respectively, * p<0.05, ** p<0.01, *** p<0.001 and **** p<0.0001).

### Correlation between neuron maturation and nano-neuro interaction

In further delving into the maturation-dependent nano-neuro interaction, we made two important observations: (i) at different DIVs, a varying fraction of neurons within the network did not interact with negatively charged PFs (**Figure 6A**, pointed with yellow arrows) and; (ii) the localized nanoparticle density varies over a wide range among the labelled neurons (**Figure 6A**, pointed with white arrows identifying low, medium and high density of PFs). We hypothesized that this graded binding of the nanostructures to neurons might be correlated with the neuron-maturation state (*i*.*e*., higher in mature neurons and lower in young neurons). To quantify this phenomenon, we employed filament tracer module of IMARIS software (OXFORD INSTRUMENTS) to extract the morphological parameters of neurons (**Figure S23**).^37^ We selected total neurite area, total neurite length and number of neurite terminals extracted using filament tracking analysis as the morphological maturation parameters to examine the correlation between neuron maturation and nanoparticle binding.^38, 39^ The image corresponding to MAP2 channel representing all the neurons in the culture was utilized for comparing the morphological parameters of labeled and unlabeled neurons, while the image corresponding to PFs channel was only utilized to spot the neurons with and without nanoparticles. We observed that the neurons tagged with negative PFs exhibited significantly higher morphological maturation parameters as compared to the neurons without PFs (**Figure 6B, C and D**), suggesting that nanoparticle localization is highly dependent on the maturation state of the neurons. Further, the density of nanoparticles localized on the neurons (measured as fluorescence intensity of PFs) exhibited strong correlation (Pearson’s r value of 0.81) with the morphological maturation parameters of neurons (**Figure 6E, F and G**). These observations suggest that the morphological maturation stage of the neurons possibly regulates the interactions between nanoparticles and neurons with higher binding on mature neurons and lower binding on maturing neurons.

**Figure 6.**
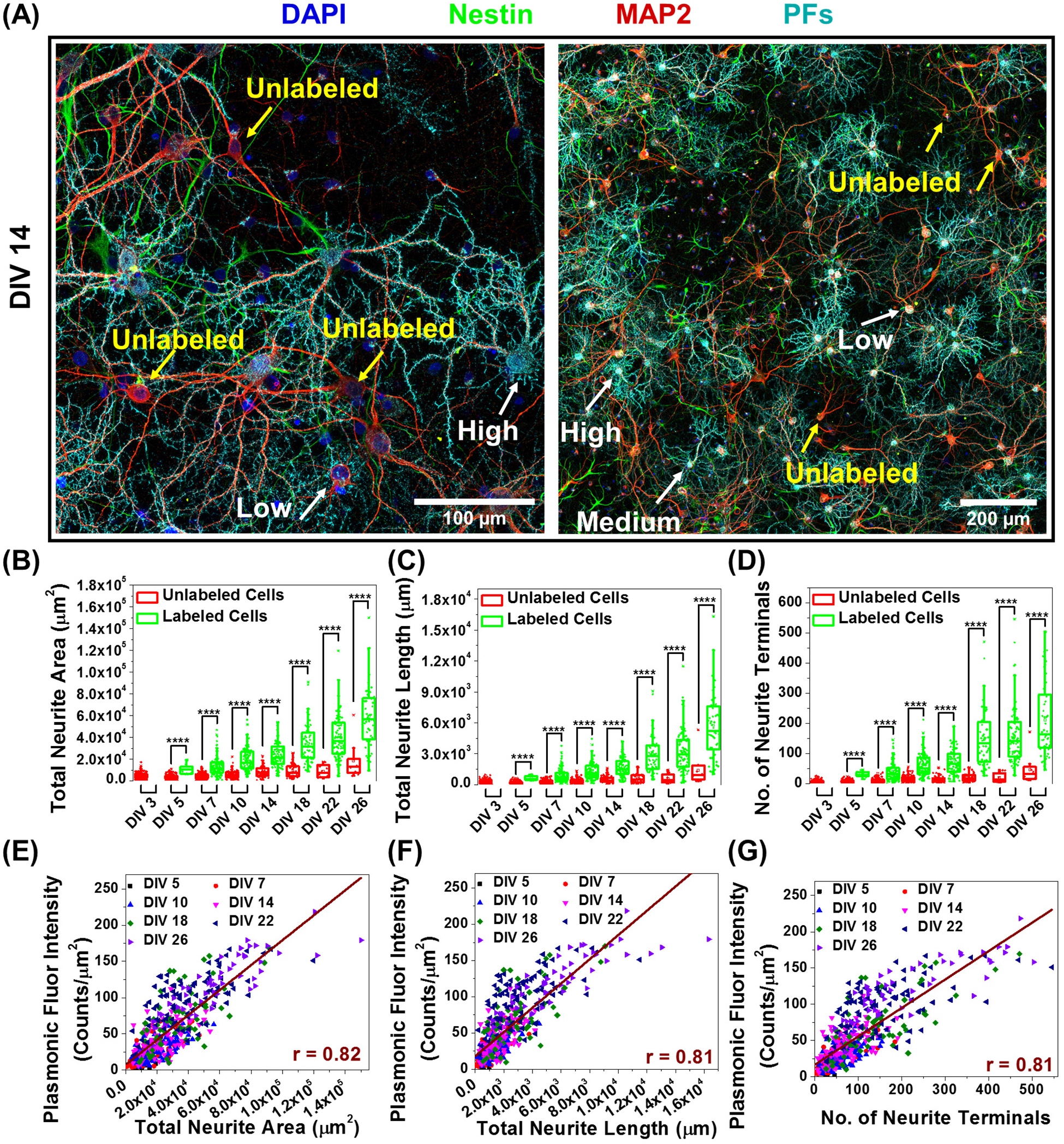
Correlation between morphological maturation parameters of neurons and the nano-neuro interaction. (A) Confocal fluorescence images of cultured hippocampal neurons after 1 hour incubation with negatively charged PFs (cyan) at DIV 14, co-stained with MAP2 (red, neuronal cell marker, specific to neuron cells), Nestin (green, progenitor cell marker, stains both neurons and glial cells) and DAPI (blue, nucleus staining). The left panel shows the fluorescence image at 20× magnification and the right panel shows the 3×3 tiled image obtained from 9 images similar to left panel. (n=2 independent experiments). Yellow arrows indicate untagged cells and white arrows indicate tagged cells with graded tagging. Whisker plot representing the morphological maturation parameters: (B) total neurite area, (C) total neurite length and, (D) no. of neurite terminals of neurons with and without nanoparticles at DIV 3, 5, 7, 10, 14, 18, 22 and 26. Morphological maturation parameters were extracted from fluorescence images using MAP2 and PF channels via filament tracking analysis. Unpaired Two-samples t-test; n = 215, 190, 134, 94, 64, 42, 15 and 10 unlabeled neuronal cells and n = 0, 5, 106, 111, 94, 98, 95 and 54 labeled neuronal cells from three 3×3 tiled images from the same culture for DIV 3, 5, 7, 10, 14, 18, 22 and 26 respectively, * p<0.05, ** p<0.01, *** p<0.001 and **** p<0.0001. Correlation between morphological maturation parameters: (E) total neurite area, (F) total neurite length and, (G) no. of neurite terminals and the fluorescence intensity of PFs bound on the neuron (which is directly related to density of PFs on the neuron). The scatter plot is presented using the data from all the tagged cells and Pearson’s correlation coefficient (r) is calculated after performing linear fitting of the concatenated data.

## Discussion

We elucidate the effect of nanoparticle binding to neurons on the electrical activity in a neuronal network as well as the effect of maturation-dependent nano-neuro interaction on the nano-neuromodulation. The negative surface charge of the nanoparticles is a necessary condition for spontaneous binding of the nanoparticles to neurons in a culture. These results are in agreement with a recent report that highlighted the importance of the surface of charge of nanoparticles in nano-neuro interactions and the relative insignificance of the size, shape, and composition of the nanostructures.^26^ It has been reported that the nano-neuro interaction depolarizes neuronal membrane potential, resulting in increased excitability and firing rate of individual neurons.^40, 41^ The increase in the excitability of the neurons results in increased probability of burst discharge instead of single firing event.^42^ Consequently, in the present study, as a result of nanoparticle binding to a fraction of neurons in the neuronal network, the excitability of only the neurons tagged with nanoparticles is expected to increase. The increased excitability of tagged neurons in turn significantly increased the bursting activity of the network. Consequently, this heterogeneous nano-neuro interaction resulted in faster, regular and smaller burst discharge as compared to slower, irregular and longer burst discharge in the absence of nanoparticles.

Neuronal maturation is a dynamic and heterogeneous process in which neuron undergoes well-defined transition in morphology, excitability and connectivity in the pathway toward fully mature phenotype.^36^ During the neuron maturation process, the dendritic length and the number of dendritic terminals of the neurons increase. Moreover, during this maturation process, the electrophysiological properties of neurons transforms from high input resistance, relatively depolarized resting membrane potential and small action potentials in the case of young neurons to low input resistance, relatively hyperpolarized resting membrane potential and large action potentials in the case of mature neurons.^43, 44^ In the current study, we have unveiled the critical role of neuronal maturation on the nano-neuro interactions. Dante et al. indicated that the neuronal spiking activity causes the spontaneous binding of the negatively charged nanoparticles to the surface of the electrically-active neurons.^26^ We have also observed the selective binding of negatively charged particles to the neurons, which is in agreement with the previously reported work. However, upon pharmacologically manipulating the electrophysiological activity of the neural network using bicuculline (BICU) and tetrodotoxin (TTX) for increasing and suppressing the spiking activity respectively, we observed no significant effect on the nano-neuro interaction (Figure S24). This suggests that electrical activity of the neurons is not the governing factor for selective binding of negatively charged nanoparticles to neurons. Although we noted a strong correlation between morphological maturity of neurons and the binding of the negatively charged nanoparticles to the neurons, the underlying electrophysiological and/or cell surfaceome factors responsible for this maturation-dependent nano-neuro interactions still remains unclear. Oostrum et. al. recently demonstrated the reorganization of neuronal surface proteins during maturation in culture, which is proteostasis-independent and this regulation affects the quantitative surface abundance of surfaceome with very few qualitative alterations.^45^ Considering the unimportance of electrical activity of neurons in the binding of negatively charged nanoparticles to mature neurons, we speculate that the selective nano-neuro interaction might be attributed to this surfaceome reorganization with maturation, which might lead to change in the surface charge of the neurons from neutral in early stages to positively-charged with maturation. It is important to note that the interaction of positively charged and negatively charged nanoparticles with the cells is still considered as a standard technique to estimate the surface charge of living as well as fixed cells,^46, 47^ which is similar to the experimental protocol employed in this work, suggesting the progressive change in the neuronal surface charge with maturation. This change in the neuronal surface charge with maturation might be responsible for the maturation-dependent graded nano-neuro interaction of negatively-charged nanoparticles. Considering that a fine control over nano-neuro interactions is critical for efficacious nano-neuromodulation, the mechanistic aspects of interaction between negatively charged particles and neuronal membrane needs to be further investigated. Regardless, this heterogeneous nano-neuro interaction elicited major implications on the optical neuromodulation of the cultured neurons. Plasmonic nanostructures have been widely investigated for neuromodulation, which either resulted in inhibition of spiking activity under continuous wave laser stimulation or increase in spiking activity via pulsed laser illumination.^12, 17^ In the case of pulsed laser, a likely mechanism for the increase in spiking activity is the photothermally induced membrane capacitance changes resulting in cell depolarization.^48, 49^ Another possible mechanism is the alteration of cell membrane properties via short thermal pulses. Potentially, short thermal pulses result in transient nanopores in the cell membrane, which in turn can increase cell membrane fluidity, thereby altering the cell potential and activating the voltage-gated ion channels.^50-52^ On the other hand, the inhibition of spiking activity in response to continuous wave lasers likely stems from the thermo-sensitive potassium channel – TREK-1.^17^ In stark contrast to the earlier observations, for the first time, we report a heterogeneous modulation in neuronal activity at network level *i*.*e*., simultaneous excitation and inhibition of electrical activity under optical stimulus. The heterogeneous optical modulation in the early stages (*i*.*e*. at DIV 14 and 18) of neuronal maturation may be attributed to heterogeneous binding of the nanostructures to neurons (∼65-70% of the neurons in the culture tagged with nanoparticles). We speculate that the channels of MEAs exhibiting partial inhibition or no change in spiking activity majorly comprise of maturing neurons, which do not respond to photothermal stimulation because of the absence or low density of photothermal nanostructures on their surface.^25^ On the other hand, the electrodes exhibiting increased spiking activity might be majorly surrounded by un-tagged young neurons. Since these untagged neurons are part of a larger neuronal network, photothermal stimulation could turn down inhibitory inputs received from maturing or matured neurons thereby resulting in an increased spiking activity of these young neurons.^25^ Similarly, there can be many other combinations of young, maturing and mature neurons that can result in the observed heterogeneous response to photothermal stimulation. In contrast, as majority of the neurons reach advanced states of maturity or are completely matured, the photothermal stimulation resulted in nearly complete inhibition of spiking activity as observed in the case of DIV 22 and 26. This observation suggests that till DIV 18, a major fraction of neurons in the network are still young or in early maturation stage while after DIV 22, majority of the neurons reached maturing or matured state. This is also evident from the fluorescence images demonstrating the binding of negatively charged PFs on the neurons at corresponding DIVs. The graded and selective binding of negatively charged nanoparticles on neurons demonstrated here opens novel avenues in minimally-invasive nanomaterials based non-genetic neuromodulation approach for treatments of neural disorders in the complex environment of large mammalian brain.

### Outlook

In summary, we unveil neuronal maturation-dependent nano-neuro interaction and nano-neuromodulation with a transformation from heterogeneous to homogeneous change in electrical activity with photothermal stimulation. We found that the nano-neuro interaction not only depends on the surface charge of the nanoparticles but also strongly correlates with the maturation stage of each individual neurons in the network, which in turn determines the homogeneity of nano-neuromodulation in a maturing neural network. Our results have broad implications in both neuroscience research as well as clinical applications. Recent advances in nanotechnology have revolutionized the field of neuroscience research by enabling various neuromodulation modalities (viz. electrical, optical, acoustic, magnetic, chemical).^1, 10, 53^ A comprehensive understanding of the factors influencing nano-neuro interaction will greatly advance our capability to seamlessly integrate nanomaterials with the nervous system and could help shape the future of neuromodulation therapy. Furthermore, we envisage that the ability to achieve precise and selective nano-neuro interaction could potentially alleviate the incessant bottleneck in the deployment of nanoprobes such as plasmonic NPs, up conversion nanoparticles, quantum dots, and nanodiamonds for both recording as well as manipulating complex neural ciruits in vitro and *in vivo*.^54-57^

Considering the critical role of nanoparticle surface charge on nano-neuro interaction, the heterogeneity observed in nano-neuromodulation, stemming from the heterogeneous binding of nanoparticles in a maturing neural network, possibly applies to wide repertoire of nano-enabled neuromodulation modalities. Moving forward, we envision that a better understanding of the cell surface proteins responsible for the maturation-dependent electrostatic state of the neurons, possibly governing the maturation-dependent nano-neuro interactions presented here, could prove as an additional tool in the nanotechnology toolkit for the development of next-generation neuromodulation modalities with unprecedented spatiotemporal resolution. In clinical applications, owing to the fact that neurogenesis persists throughout aging in human hippocampus,^24, 25, 36^ the maturation-dependent graded and selective nano-neuro interaction offers an additional handle in developing nano-neuromedicine for addressing neurological disorders. Moreover, the change in firing pattern of the neural network stemming from nano-neuro interaction could serve as a non-invasive treatment for diseases that are characterized by erratic electrical activity in parts of the brain, such as epilepsies and seizures.^58, 59^ Collectively, our findings facilitate the development of new nanotechnologies for nano-neuro interface, which may be broadly applicable to both understanding neural pathways as well as minimally-invasive nano-enabled drug-like administrable neurotherapeutics.

## Methods

### Cell culture

All procedures have been approved by the Institutional Animal Care and Use Committee (IACUC) at Washington University in St. Louis. The hippocampal tissues were manually isolated from day E18 embryos of pregnant Sprague Dawley rat brains (Charles River, USA) in Hibernate EB medium (HEB, BrainBits, USA) as previously described.^23^ The isolated tissues were incubated in cell dissociation solution comprising of 6 mg papain (P4762, Sigma, USA) in 3 ml of Hibernate E-Ca (HE-Ca, BrainBits, USA) for 10 minutes at 30°C. Subsequently, the tissues were mechanically dissociated via trituration with fire-polished Pasteur pipette after replacing cell dissociation solution with HEB medium to obtain single cell suspension. The resultant cell suspension was centrifuged at 200xg for 1 minute and the supernatant was decanted, and the pellets were resuspended in NbActiv4 medium (BrainBits, USA). Prior to plating the cells, the substrates were coated with poly(ethyleneimine) solution (0.1 % in water, P3143, Sigma, USA) for 30 minutes followed by rinsing with water, air drying and sterilization under UV light exposure for 1 hour. Subsequently, the substrates were further treated with laminin solution (20 μg ml^−1^ in NbActiv4 medium, L2020, Sigma, USA) for 30 minutes to promote cell adhesion and neurite outgrowth.^60^ After decanting the excess laminin solution from the substrates, the cells were plated onto glass bottom petri dishes (35 mm Glass bottom dish with 14 mm micro-well #1 cover glass, D35-14-1-N, Cellvis, CA, USA) at a density of 120 – 160 cells/mm^2^ for use in microscopy experiments and onto microelectrode array (MEA, Multichannel Systems, Germany) at a density of 500 – 1000 cells/mm^2^ for electrophysiology measurements. The neurons were maintained in a humidified incubator with 5% CO_2_ and 37 ºC condition. At DIV 3, half of the culture medium (NbActiv4) was replaced with fresh culture medium and subsequently replaced regularly every 7 days in case of glass bottom petri dishes and every 2 days in the case of MEAs.

### Synthesis of positively-charged PFs

The PF-650 with IR-650 as molecular fluorophore and Au@Ag nanocuboids as plasmonic core were generously provided by Auragent Bioscience, MO, USA. The PFs were synthesized according to a procedure we recently reported.^27^ Owing to the presence of BSA on the surface of the PFs, the PFs are inherently negatively charged under physiological pH. To realize positively charged PFs, the surface of these negatively charged PFs were coated with cationic polyelectrolyte, poly (allylamine hydrochloride) (PAH, 43092, Alfa Aesar, USA), via electrostatic interaction. Briefly, 10 ml of PFs (O.D. ∼ 2) was washed 3 times using alkaline nanopure water (pH = 10) via centrifugation at 6000 rpm to remove excess salt present in the storage buffer of PFs and re-dispersed in 10 ml of nanopure water (pH = 10). Subsequently, the purified PFs were added dropwise to 10 ml of PAH solution (0.2% W/V in water, pH adjusted to 10 using 1M NaOH) under vigorous stirring and sonicated for 1 hour at room temperature under dark condition. Finally, PAH-coated PFs were washed with nanopure deionized (DI) water (resistivity >18.2 MΩ.cm) twice by centrifugation at 6000 rpm and re-dispersed in DI water for further use.

### Synthesis of negatively charged AuNRs

The localized surface plasmon resonance (LSPR) wavelength of the AuNRs can be easily tuned over a wide range by varying their aspect ratio.^61-63^ Considering the wavelength of NIR light source (808 nm) utilized for photothermal modulation in the present work, the AuNRs with LSPR wavelength of 820 nm were synthesized via previously reported seed-mediated approach.^61, 62, 64^ Briefly, the gold seed solution was first prepared by adding 0.6 ml of ice-cold 10mM NaBH_4_ solution (71321, Sigma, USA) into a magnetically stirred (800 rpm) solution comprising of 0.25 ml of 10 mM HAuCl_4_ (520918, Sigma, USA) and 9.75 ml of 0.1 M hexadecyltrimethylammonium bromide (CTAB) (H5882, Sigma, USA) at room temperature for 10 min. Consequently, the solution color changed from orange to brown, indicating the Au seed formation. Subsequently, the growth solution was prepared by sequential addition of 2 ml 10 mM HAuCl_4_ aqueous solution, 38 ml 0.1 M CTAB solution, 0.4 ml 10 mM AgNO_3_ (204390, Sigma, USA) and 0.22 ml 0.1 M ascorbic acid (A92902, Sigma, USA) followed by gentle homogenization via inversion, rendering growth solution color change from orange to colorless. Finally, 48 μl of the freshly prepared gold seed solution was added to the growth solution, mixed via inversion and left undisturbed in the dark at room temperature for 24 hours. The AuNRs were collected via centrifugation at 9000 rpm for 30 min to remove the supernatant and re-dispersed in DI water for further use.

Owing to the presence of CTAB on the surface of AuNRs, the AuNRs are inherently positively charged. To realize negatively-charged AuNRs, the positively-charged AuNRs were coated with anionic polyelectrolyte, poly (sodium 4-styrenesulfonate) (PSS, 434574, Sigma, USA), via electrostatic interaction. Briefly, 10 ml of AuNRs (O.D. ∼ 2) was washed once with DI water via centrifugation at 9000 rpm to remove excess CTAB and re-dispersed in 10 ml of nanopure deionized (DI) water (resistivity >18.2 MΩ·cm). Subsequently, the purified AuNRs were added dropwise to 10 ml of PSS solution (0.5% W/V in water) under vigorous stirring and sonicated for 1 hour at room temperature. Finally, the PSS-coated AuNRs were washed with DI water twice by centrifugation at 9000 rpm and re-dispersed in DI water for further use.

### Material Characterization

Transmission electron microscopy (TEM) micrographs were acquired using JEOL JEM-2100F field emission microscope. A drop of plasmonic nanostructure aqueous dispersion was casted onto the copper grids (Carbon Type-B, 200 mesh, Ted Pella, USA). The extinction spectra of plasmonic nanostructures were acquired using shimadzu UV-1800 spectrophotometer. The zeta potential measurements were performed using Malvern Zetasizer (Nano ZS). Large area fluorescence mappings were obtained using LI-COR Odyssey CLx imaging system.

### Neuron electrophysiology experiments

#### Neural recording

Extracellular electrophysiological recordings from primary cultured hippocampal neurons were performed using 60-channel TiN microelectrode arrays (60MEA200/30iR-Ti-gr, MultiChannel Systems, electrode diameter 30 μm, electrode spacing 200 μm, 8 × 8 electrode grid, 59 electrodes, 500 nm thickness of Si_3_N_4_ insulator). The extracellular recordings of the spontaneous network activity were acquired simultaneously from all the 59 electrodes utilizing an *in vitro* MEA recording system (MEA2100-Mini-System, Multichannel systems, gain 1100, bandwidth 10-8 kHz, sampling frequency 25 kHz). The electrodes were maintained at 37°C and 5% CO_2_ atmosphere via a climate chamber (MEA2100-CO2-C, MultiChannel Systems) during electrophysiological recordings. The recording of the neuronal activity was performed 20 min after placing the MEA in the recording system equipped with climate chamber. The recorded raw voltage traces were filtered with a 200 Hz digital high pass filter (Butterworth, second order), and the spikes were detected by defining the threshold level as six times of the standard deviation of background noise using a software provided by the vendor (MC Rack, MultiChannel Systems). The network bursts were detected utilizing MaxInterval algorithm,^65^ available in the software (MC Rack, MultiChannel Systems) by defining minimum number of spikes in a burst as 4, maximum interspike interval to start the burst as 100 ms, maximum interspike interval to end the burst as 500 ms, minimum interspike interval between two bursts as 500 ms and minimum burst duration as 20 ms. Collected data were processed using custom built MATLAB (MathWorks) script.

#### Nano-neuro interaction

To assess the effect of nano-neuro interaction on the electrophysiology of the neurons, the neuronal network activity of the cultured hippocampal neurons at DIV 14 was recorded for 10 minutes prior to nanoparticle administration. Subsequently, the negatively-charged AuNRs dispersed in NbActiv4 medium was added to the culture medium at a final concentration of O.D. ∼ 0.5. After 1 hour incubation with nanoparticles, the activity of the neural network was recorded for 10 mins. The recording channels with average firing rate greater than 0.1 spikes/sec were selected as active channels and utilized for further neural activity analysis. Subsequently, the effect of nanoparticle binding on the neuronal activity was analyzed utilizing following four main parameters: (i) mean spike rate, calculated as average firing rate over entire recording duration; (ii) mean burst rate, calculated as average number of bursts per minute; (iii) burst duration, calculated as average duration of burst events; and (iv) mean spikes per burst, calculated as average number of spikes during burst events. All statistical difference between two groups were analyzed using unpaired one-tailed t-test with 5% one-sided significance level.

#### Maturation-dependent nano-neuromodulation

To investigate the maturation-dependent nano-neuromodulation, the photothermal modulation of neuronal network activity of the cultured hippocampal neurons on a MEA chip at DIV 14, 18, 22 and 26 was performed. To avoid any interference, the nano-neuro interaction might have on maturation of the neurons, separate MEAs were employed for neuromodulation experiments at different DIV. The negatively charged AuNRs dispersed in NbActiv4 medium were added to the neuron culture on specific DIV at a final concentration of O.D. ∼ 0.5 and incubated for 1 hour in the incubator maintained at 37°C and 5% CO_2_. To minimize the free AuNRs in the culture, prior to neuromodulation experiments, the AuNR treated cultures were gently washed three times with NbActiv4 medium by replacing 75% of the medium with fresh medium followed by gentle swirling. Subsequently, the MEAs were placed in the incubator for yet another hour for stabilization. A fiber optic coupled NIR laser diode module (808 nm, continuous wave, 2 W, Power technologies inc.) was utilized as a light source for photothermal neuromodulation and the collimator present at the end of the optical fiber provides a means to tune the laser beam spot size and power density by controlling its distance from the MEAs. A typical photothermal neuromodulation experiment lasts for 480 seconds, and the AuNR treated neurons on MEAs were illuminated with NIR laser at a power density of 14 mW/mm^2^ for 10, 20, 30 and 60 seconds while simultaneously recording the neuronal network activity during the entire time period of the experiment. A mechanical shutter was employed to control the laser on and off period. The recording channels with average firing rate greater than 0.1 spikes/sec were selected as active channels and utilized for further neural activity analysis. peristimulus time histogram and raster plots were used to analyze the photothermal neuromodulation with NIR irradiation as a stimulus. The spike rate change under NIR stimulus was calculated by the following equation: ΔR/R (%) = [R(ON) - R(OFF)] × 100/ R(OFF), where R(OFF) and R(ON) represents the mean spike rate before and after the onset of NIR stimulus, respectively. R(OFF) includes the 60 second window just before the onset of the stimulus and R(ON) includes the entire stimulus period. The channels exhibiting less than 10% change in the electrical activity in response to NIR stimulus were categorized as channels with no effect to photothermal modulation.

#### Neural network activity alteration

To assess the efficacy of pharmacological agents in altering the electrophysiological activity of the neuronal network, the neuronal network activity of the cultured hippocampal neurons on MEAs at DIV 14 and 26 was recorded for 5 minutes prior to subjecting the network to the specific pharmacological agent. Subsequently, either tetrodotoxin (TTX, ab120055, Abcam, USA) or bicuculline (BICU, 14340, Sigma, USA) dissolved in NbActiv4 medium was introduced into the cultured neurons at a final concentration of 1 μM and 30 μM, respectively. After 15 minutes of incubation with the pharmacological agents, the activity of the neuronal network was recorded for 5 minutes. The recording channels with average firing rate greater than 0.1 spikes/sec were selected as active channels and utilized for further neural activity analysis. Subsequently, the efficacy of TTX and BICU to suppress or increase the neuronal network activity, respectively, was analyzed utilizing mean spike rate, calculated as average firing rate over entire recording duration, before and after administration of the pharmacological agents.

### Confocal fluorescent microscopy experiments

#### Assessing nanoparticle surface charge dependent nano-neuro interaction

PFs with different surface charges dispersed in Nbactiv4 medium were administered to the cultured neurons on DIV 3, 5, 7, 10, 14, 18, 22 and 26 at a final concentration of O.D. ∼ 0.5 and incubated for 1 hour at 37°C and 5% CO_2_. Subsequently, the cells were washed once with 1X phosphate buffered saline (PBS) and were fixed with 4% (W/V) paraformaldehyde (PFA, Sigma, USA) solution in PBS at room temperature for 30 min, followed by washing 3 times with PBS. Finally, the nucleus was stained with DAPI (Sigma, USA) at a concentration of 300 nM in PBS for 5 minutes, followed by washing 3 times with 1X PBS. Cells were visualized under inverted confocal fluorescent microscope (Lionheart FX Automated Microscope, BioTek, USA).

#### Assessing viability of cells not tagged with negatively charged PFs

Negatively charged PFs dispersed in Nbactiv4 medium were added to the cultured neurons on DIV 14 at a final concentration of O.D. ∼ 0.5 and let to incubate for 1 hour at 37°C and 5% CO_2_. Subsequently, the cells were stained with a LIVE/DEAD cell viability assay kit (L3224, Thermo Fisher Scientific, USA), followed by fixation with PFA and nuclei staining with DAPI as discussed previously. Cells were visualized under inverted confocal fluorescent microscope (Zeiss LSM 880 Airyscan Two-Photon Confocal Microscope, Carl Zeiss AG, Germany). The viability of the neurons both labeled and unlabeled with negatively charged PFs was assessed by analyzing the presence of green-fluorescent calcein-AM stain corresponding to live neurons both with and without PF co-localization.

#### Immunostaining

Negatively charged PFs dispersed in Nbactiv4 medium were added to the neuron culture on DIV 3, 5, 7, 10, 14, 18, 22 and 26 at a final concentration of O.D. ∼ 0.5 and let to incubate for 1 hour at 37°C and 5% CO_2_. The cells were washed with PBS once, fixed with PFA and permeabilized with 0.5% Triton X-100 in PBS for 10 minutes at room temperature, followed by washing with PBS 3 times. To avoid the non-specific binding of antibodies, the cells were blocked with blocking solution comprising of 6% bovine serum albumin (BSA, Sigma, USA) in PBS for 30 minutes and washed once with 0.05% Tween-20 (Sigma, USA) in PBS. The cells were incubated with primary antibodies, mouse anti-MAP2 (2 μg/ml, monoclonal, MA5-12826, Thermo Fisher Scientific, USA) and goat anti-Nestin (10 μg/ml, polyclonal, PA5-47378, Thermo Fisher Scientific, USA), diluted in blocking solution. After 3 hour incubation at room temperature, the cells were washed with PBS three times and incubated with secondary antibodies, Alexa Fluor 568 labelled Donkey anti-Mouse (2 μg/ml, A10037, Thermo Fisher Scientific, USA) and Alexa Fluor Plus 488 labelled Donkey anti-Goat (10 μg/ml, A32814, Thermo Fisher Scientific, USA), diluted in blocking solution for 1 hour at room temperature. After washing with PBS three times, the nucleus was stained using DAPI as described previously. Cells were visualized under inverted confocal fluorescent microscope (Zeiss LSM 880 Airyscan Two-Photon Confocal Microscope, Carl Zeiss AG, Germany). The imaging conditions were kept constant for all the samples in order to compare the change in fluorescence intensity of PFs with maturation.

#### Confocal fluorescence imaging of neural network activity alterations

To investigate the role of neuronal network activity on the nano-neuro interaction, the binding of negatively-charged nanoparticles was assessed after pharmacologically altering the electrical activity of the cultured hippocampal neurons. After incubating the cultured neurons with 1 μM TTX or 30 μM BICU on DIV 14 and 26 for 15 minutes, the negatively charged PFs were added to the neuron culture at a final concentration of O.D. ∼ 0.5 in the presence of pharmacological agents and incubated for 1 hour at 37°C and 5% CO_2_. The cells were fixed, stained with primary and secondary antibodies, and analyzed using confocal microscopy following the same protocol as discussed in the previous section.

#### Confocal fluorescence image analysis

The confocal fluorescence images were analyzed using filament tracking module of IMARIS software (OXFORD INSTRUMENTS). The channel corresponding to MAP2, which is a neuronal cell marker, from the images were employed to extract the morphological parameters (viz. filament length, filament area and number of filament terminals) of neurons. First, the starting points (soma) of the neurons were detected by adjusting the starting point threshold and all detected cells were double checked manually after auto-detection and modified if necessary to append missed neurons or remove extra starting points. Subsequently, the threshold of the seeding points was adjusted so as to trace all the neuronal processes. Care was taken to avoid the tracing of background noise. Finally, the filament tracking was performed using the filament-tracking algorithm provided in the IMARIS software. All detected filaments were double-checked manually after automatic tracking and the thresholds were readjusted manually if necessary. Additionally, the channel corresponding to PFs in the images were utilized to identify the neurons, which are selectively targeted by negatively charged PFs. We utilized the filament tracking analysis module in the IMARIS software to measure the mean fluorescence intensity of the PFs as a surrogate to the nanoparticle localization density for each targeted neuron. Subsequently, the correlation between maturation and nano-neuro interaction was analyzed utilizing following four main parameters: (i) total neurite area, calculated as total area of filaments associated with individual neuron; (ii) total neurite length, calculated as total area of filaments associated with individual neuron; (iii) number of neurite terminals, calculated as total number of terminals in a fully traced neuron after filament tracking; and (iv) fluorescence intensity of plasmonic fluors, calculated as fluorescence intensity of PFs per unit area of neurons. Care was taken to only include those cells in the analysis whose filaments are not extended to the edge of the image volume. All statistical difference between two groups were analyzed using unpaired one-tailed t-test with 5% one-sided significance level.

### Scanning electron microscopy

Negatively charged PFs dispersed in Nbactiv4 medium were added to the cultured neurons on DIV 14 ad 26 at a final concentration of O.D. ∼ 0.5 and let to incubate for 1 hour at 37°C and 5% CO_2_. Subsequently, the cells were washed once with PBS and fixed with PFA overnight at room temperature. The cells were dehydrated with ethanol and vacuum dried before being sputter coated with 10 nm of gold metal. Scanning electron micrographs (SEM) were acquired using a JEOL JSM-7001 LVF Field Emission scanning electron microscope.

## Acknowledgements

The authors acknowledge support from Air Force Office of Scientific Research (#FA95501910394 (SS and BR)). The authors thank Institute of Materials Science and Engineering at Washington University for providing access to electron microscopy facilities. The authors also thank Dr. Jai Rudra for providing access to Lionheart FX Automated Microscope. Confocal fluorescence imaging using Zeiss LSM 880 Airyscan Confocal Microscope and subsequent image analysis using IMARIS were performed in part through the use of Washington University Center for Cellular Imaging (WUCCI) supported by Washington University School of Medicine, The Children’s Discovery Institute of Washington University and St. Louis Children’s Hospital (CDI-CORE-2015-505 and CDI-CORE-2019-813) and the Foundation for Barnes-Jewish Hospital (3770 and 4642). Confocal data was generated on a Zeiss LSM 880 Airyscan Confocal Microscope which was purchased with support from the Office of Research Infrastructure Programs (ORIP), a part of the NIH Office of the Director under grant OD021629.

## Author Information

### Authors and Affiliations

**Department of Mechanical Engineering and Materials Science, and Institute of Materials Science and Engineering, Washington University in St. Louis, St. Louis, MO, 63130, USA**

Prashant Gupta, Priya Rathi, Rohit Gupta, Harsh Baldi, Avishek Debnath, Hamed Gholami Derami & Srikanth Singamaneni

**Department of Biomedical Engineering, Washington University in St. Louis, St. Louis, MO, 63130, USA**

Quentin Coquerel & Baranidharan Raman

### Contributions

S.S., B.R., P.G. and H.G.D. conceived the project. S.S., B.R., P.G., P.R. and Q.C. designed the experiments. P.G., P.R. and H.B. performed the primary neuron cell culture for all the experiments. P.G. and P.R. performed all neuron electrophysiology and fluorescence imaging experiments. R.G synthesized the positively charged plasmonic fluors. P.G. performed the TEM imaging. P.G. and H.B. prepared samples for SEM of nanoparticles labelled neuron samples. A.D. performed the SEM imaging. P.G. performed all the neuron electrophysiology data analysis and the confocal fluorescence image analysis. Q.C. provided support and input on all experiments. S.S. and B.R. directed the research. P.G. S.S. and B.R. co-wrote the paper. All authors reviewed and commented on the manuscript.

## Corresponding Authors

Correspondence to <u>Srikanth Singamaneni</u> or <u>Baranidharan Raman</u>.

## Ethics Declaration

### Competing Interests

The authors declare the following competing financial interests: S.S. is one of the inventors on a pending patent related to plasmonic fluor technology and the technology has been licensed by the Office of Technology Management at Washington University in St Louis to Auragent Bioscience LLC, which is developing plasmonic fluor products. S.S. is one of the co-founders and shareholders of Auragent Bioscience LLC. These potential conflicts of interest have been disclosed and are being managed by Washington University in St Louis.

## Supporting Information

Supplementary figures 1 – 24.

