## Supplemental for "Neuronal Maturation-dependent Nano-Neuro Interaction and Modulation"

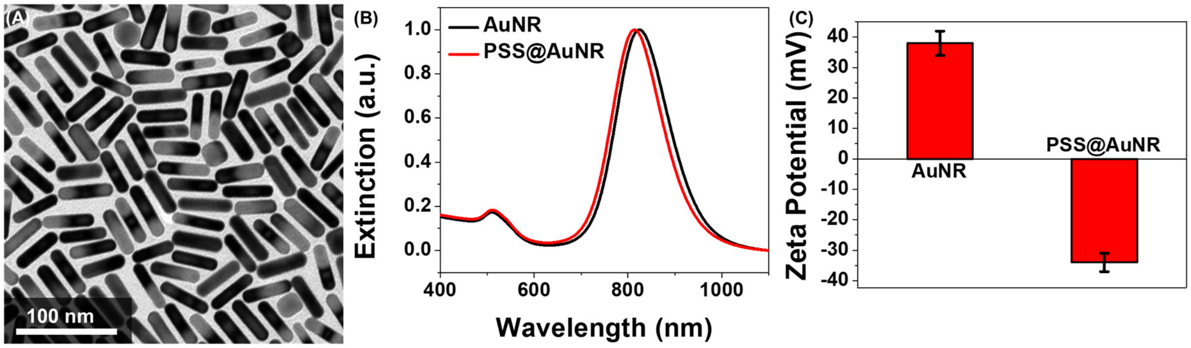


**Figure S1.** (A) The TEM micrograph of AuNR utilized for specific targeting of neurons to achieve neuromodulation. (B) Extinction spectra and (C) Corresponding zeta potential of AuNR and PSS@AuNR (negatively charged AuNRs).

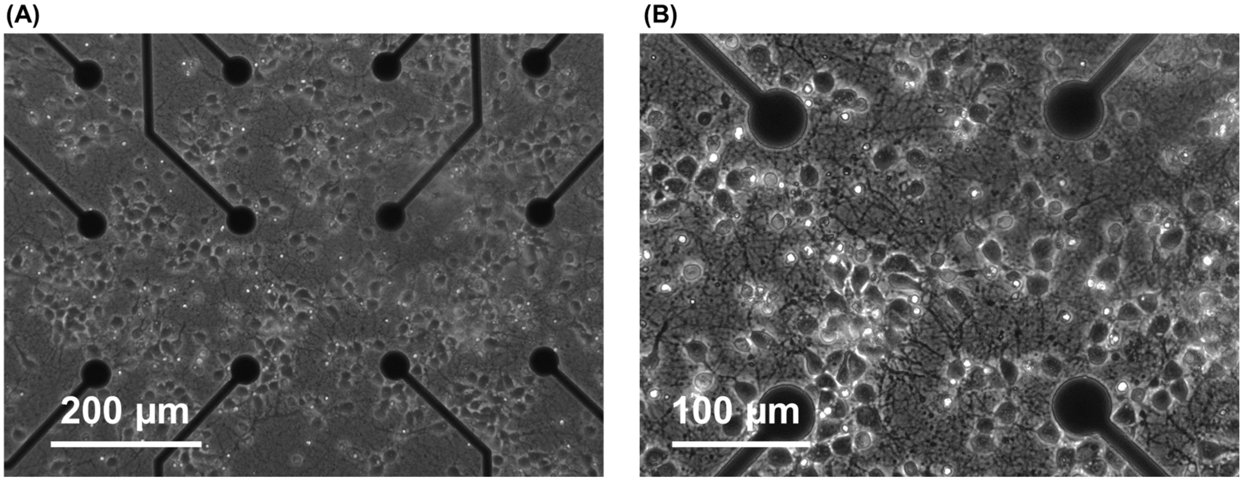


**Figure S2.** (A) Low (10X) and (B) high (20X) magnification phase contrast image of the hippocampal neurons cultured on PEI-laminin coated MEA with cell density of 1000 cells $mm^{-2}$.

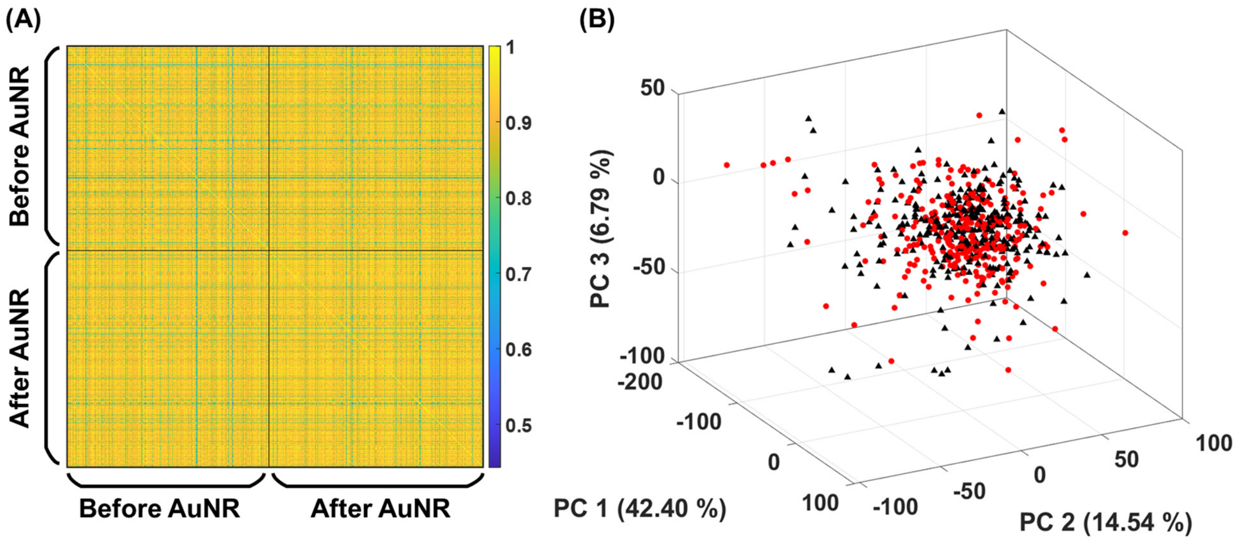


**Figure S3.** (A) correlation map and (B) Principal component analysis (first three maximum principal components, Black triangles represent spike shape before AuNR binding and red circles represent spike shape after AuNR binding) demonstrating that spike shape remains unaltered after binding of negatively charged AuNRs on the neurons. Only first 300 spikes for both conditions are presented here.

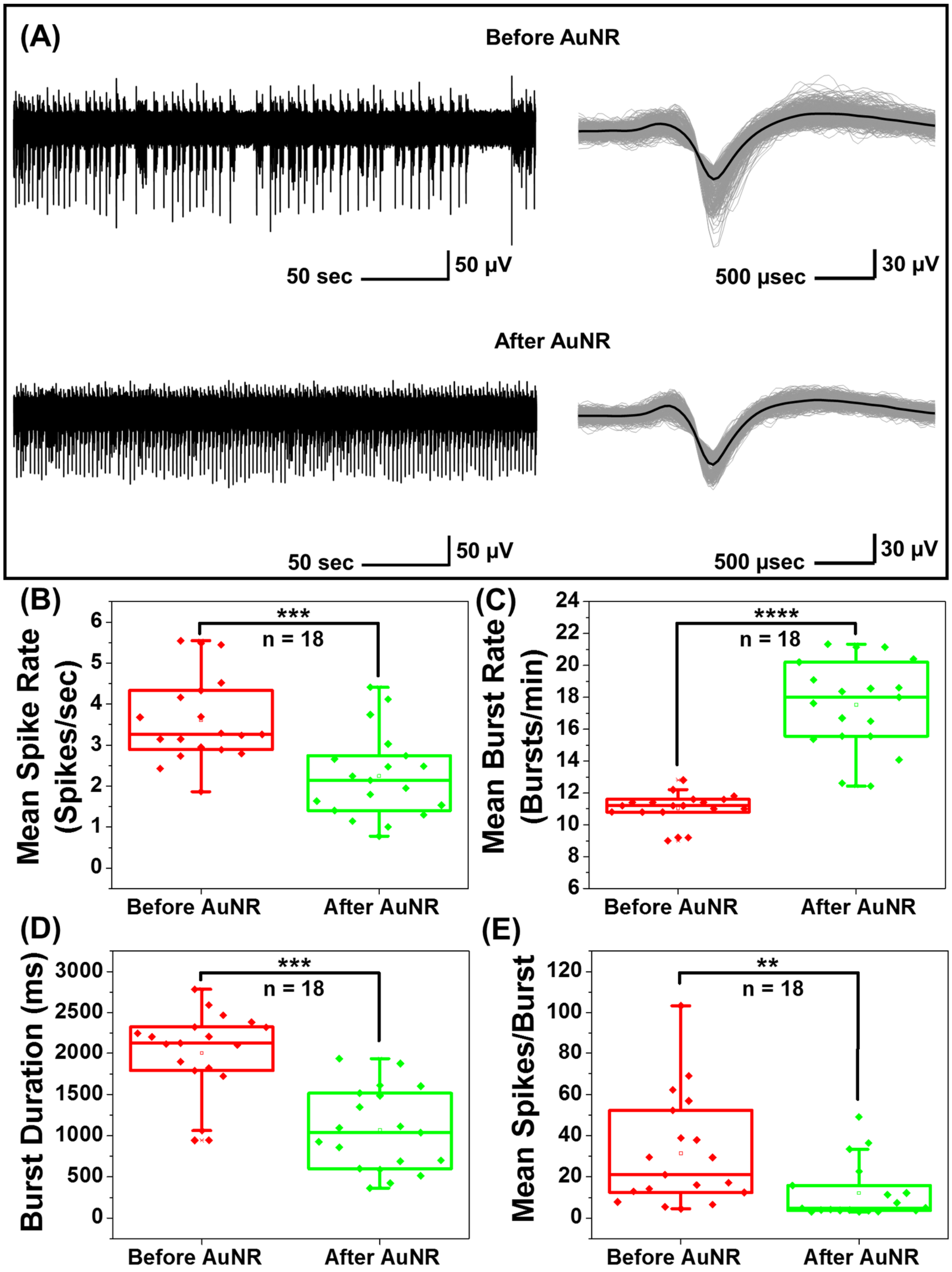


**Figure S4.** (A) A single trace of spike recording before and after neurons were incubated with negatively charged AuNR, and corresponding overlaid spike waveform of hippocampal neurons before and after AuNR labeling. Panel on the top shows the spike cutouts before the application of AuNRs and panel on the bottom shows the spike cutouts after the AuNR binding. Spikes from 5-minute recording with at least 900 spikes in each set. Black curve shows the mean value for each set. The traces are representative ones from a total of 18 active channels measured from primary cultured hippocampal neurons cultured on a microelectrode array (MEA). The experiment was repeated three times independently with similar results. Effect of AuNR localization on neuron membrane on the (B) mean spike rate (Unpaired Two-samples t-test; $p= 0.0002$, $n=18$, * $p<0.05$, ** $p<0.01$, *** $p<0.001$ and **** $p<0.0001$), (C) mean burst rate (Unpaired Two-samples t-test; $p< 0.0001$, $n=18$, * $p<0.05$, ** $p<0.01$, *** $p<0.001$ and **** $p<0.0001$), (D) burst duration (Unpaired Two-samples t-test; $p< 0.0001$, $n=18$, * $p<0.05$, ** $p<0.01$, *** $p<0.001$ and **** $p<0.0001$) and (E) mean spikes per burst (Unpaired Two-samples t-test; $p= 0.0006$, $n=18$, * $p<0.05$, ** $p<0.01$, *** $p<0.001$ and **** $p<0.0001$) of cultured neurons.

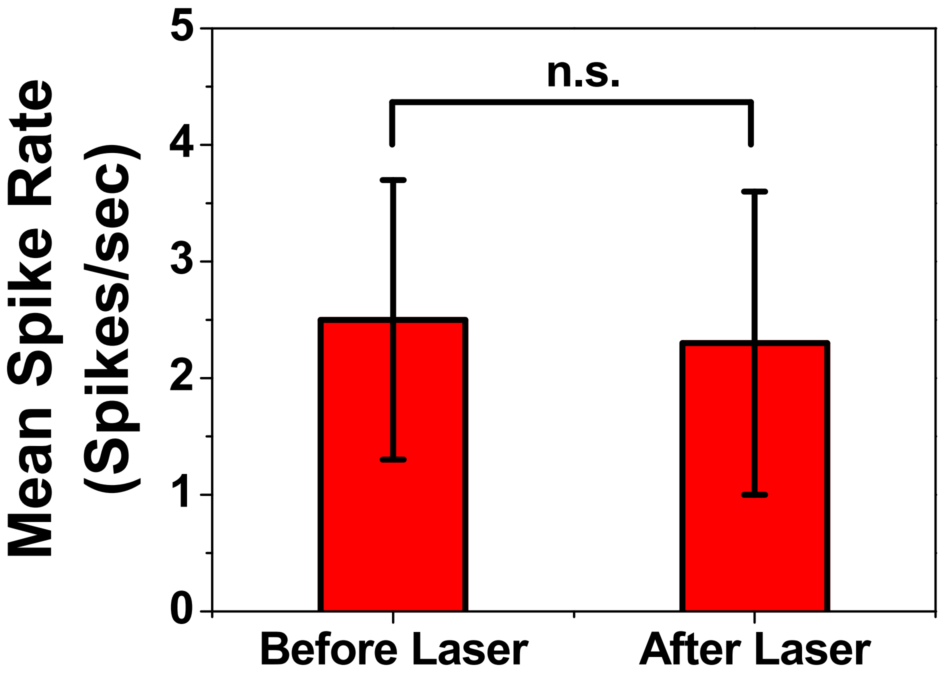


**Figure S5.** Mean spike rates of AuNR localized neurons before and after NIR irradiation (Data presented here corresponds to neuronal activities collected at DIV 14 and all the channels were considered, Unpaired Two-samples t-test; p= 0.5315, n=31).

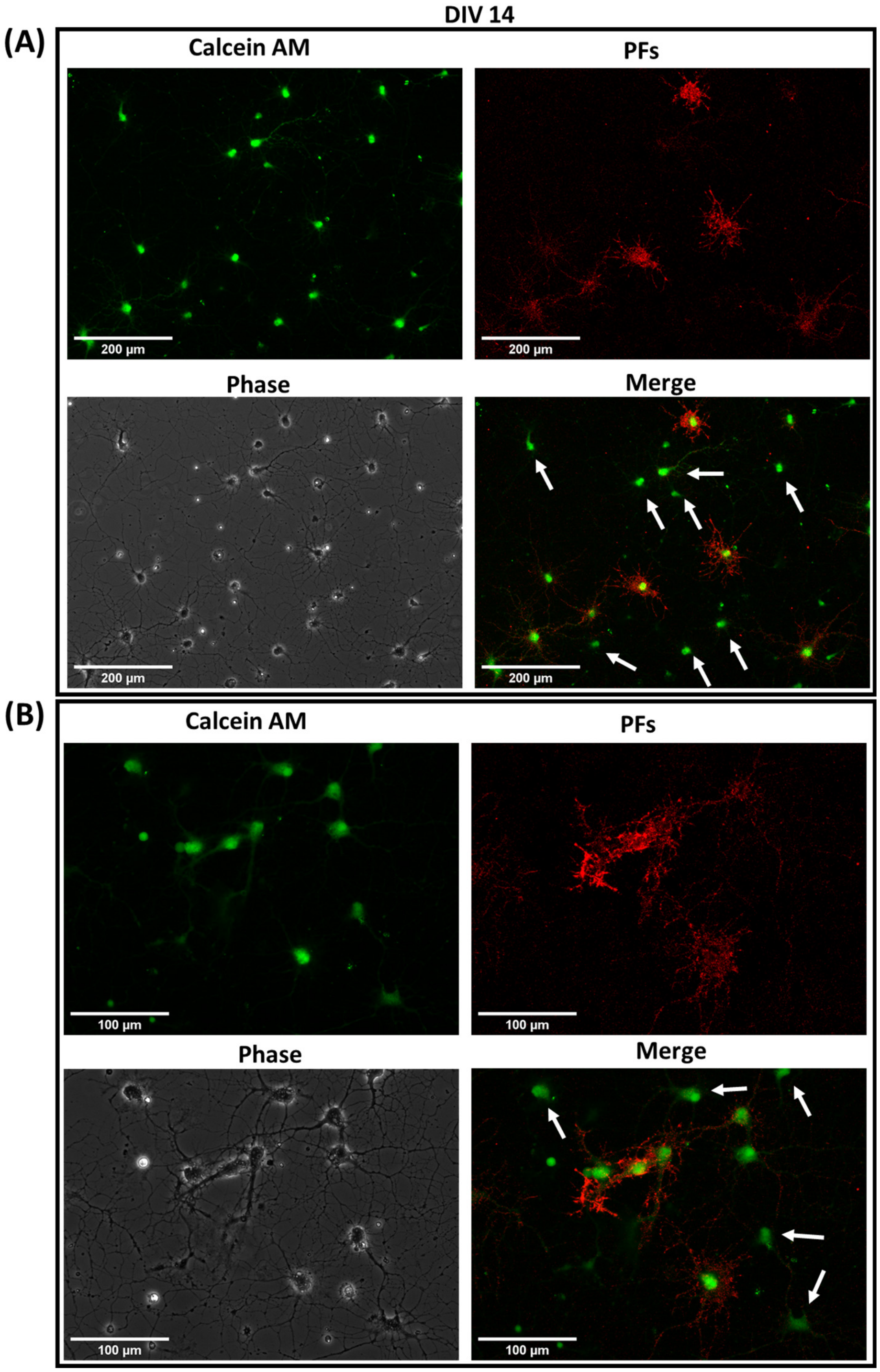


**Figure S6.** (A) Low and (B) high magnification confocal fluorescence images of cultured hippocampal neurons after 1 hour incubation with negatively charged PFs at DIV 14 (red), co-stained with calcein AM (green) for live cell staining. The white arrows indicate live cells that are not tagged with negatively charged PFs. (n=2 independent experiments)

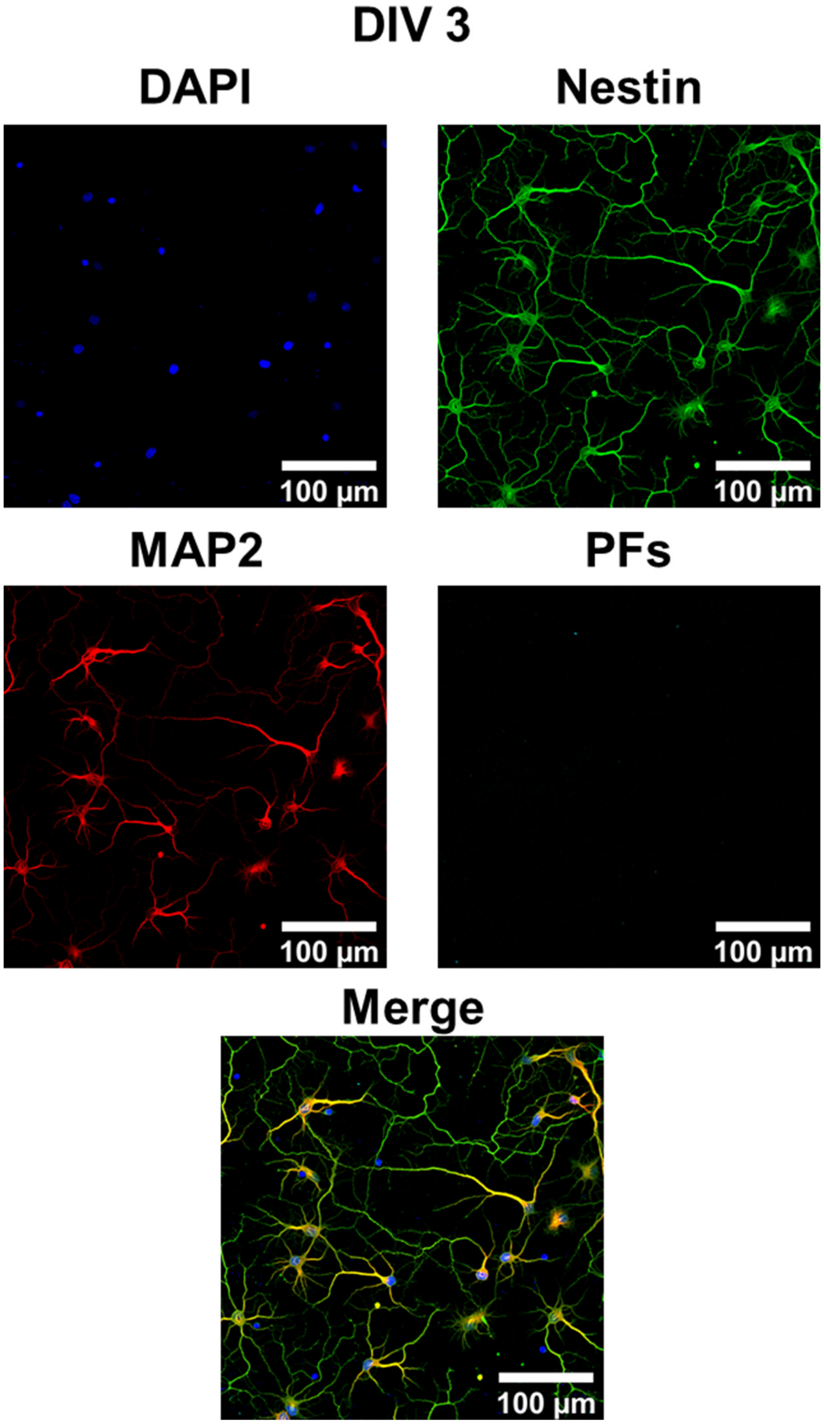


**Figure S7.** Individual channels of 20X magnification confocal fluorescence images (only merged image presented in the main text) of cultured hippocampal neurons after 1 hour incubation with negatively charged PFs (cyan) at DIV 3, co-stained with MAP2 (red, neuronal cell marker, specific to neuron cells), Nestin (green, progenitor cell marker, stains both neurons and glial cells) and DAPI (blue, nucleus staining).

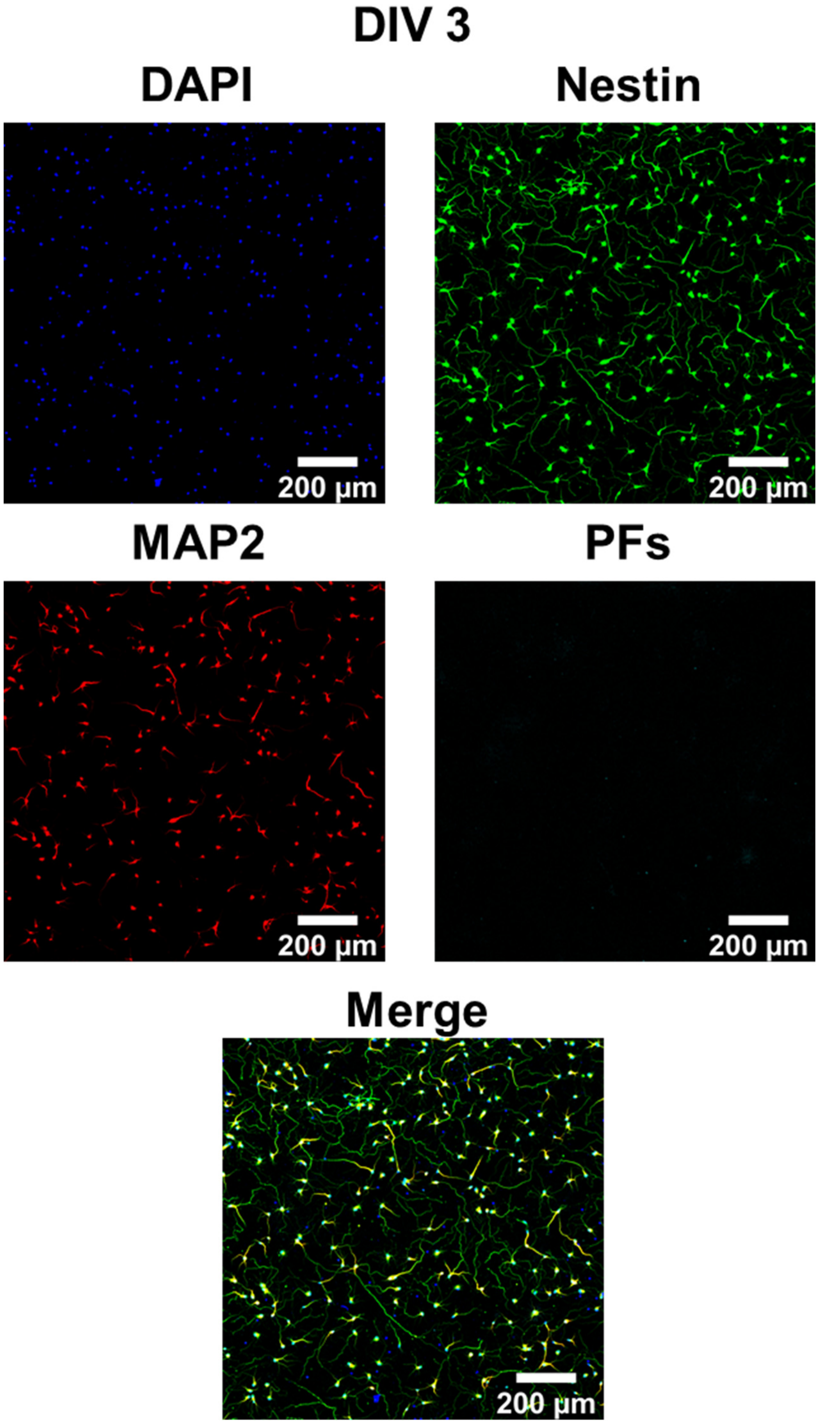


**Figure S8.** Individual channels of 3X3 tiled - 20X magnification confocal fluorescence images (only merged image presented in the main text) of cultured hippocampal neurons after 1 hour incubation with negatively charged PFs (cyan) at DIV 3, co-stained with MAP2 (red, neuronal cell marker, specific to neuron cells), Nestin (green, progenitor cell marker, stains both neurons and glial cells) and DAPI (blue, nucleus staining).

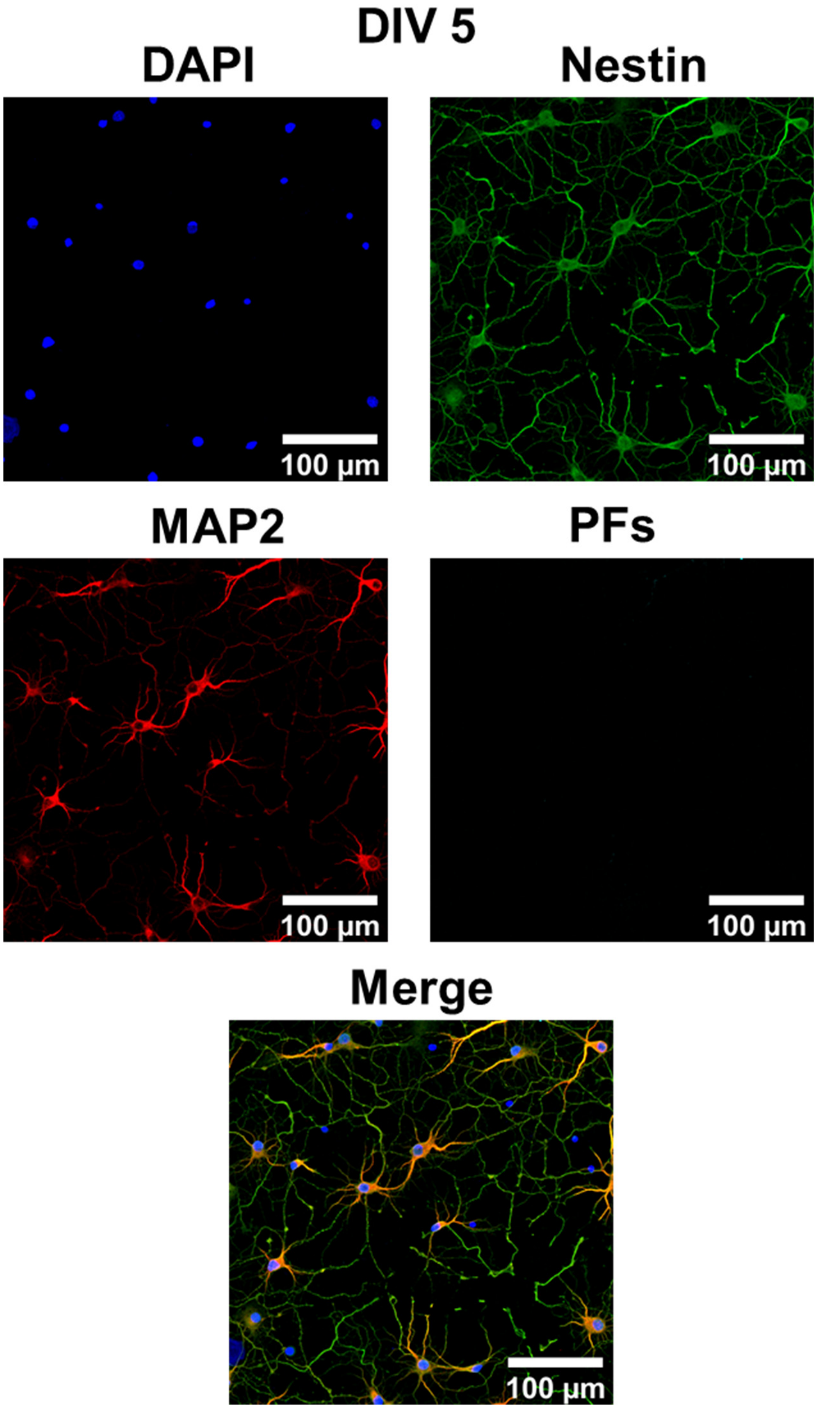


**Figure S9.** Individual channels of 20X magnification confocal fluorescence images (only merged image presented in the main text) of cultured hippocampal neurons after 1 hour incubation with negatively charged PFs (cyan) at DIV 5, co-stained with MAP2 (red, neuronal cell marker, specific to neuron cells), Nestin (green, progenitor cell marker, stains both neurons and glial cells) and DAPI (blue, nucleus staining).

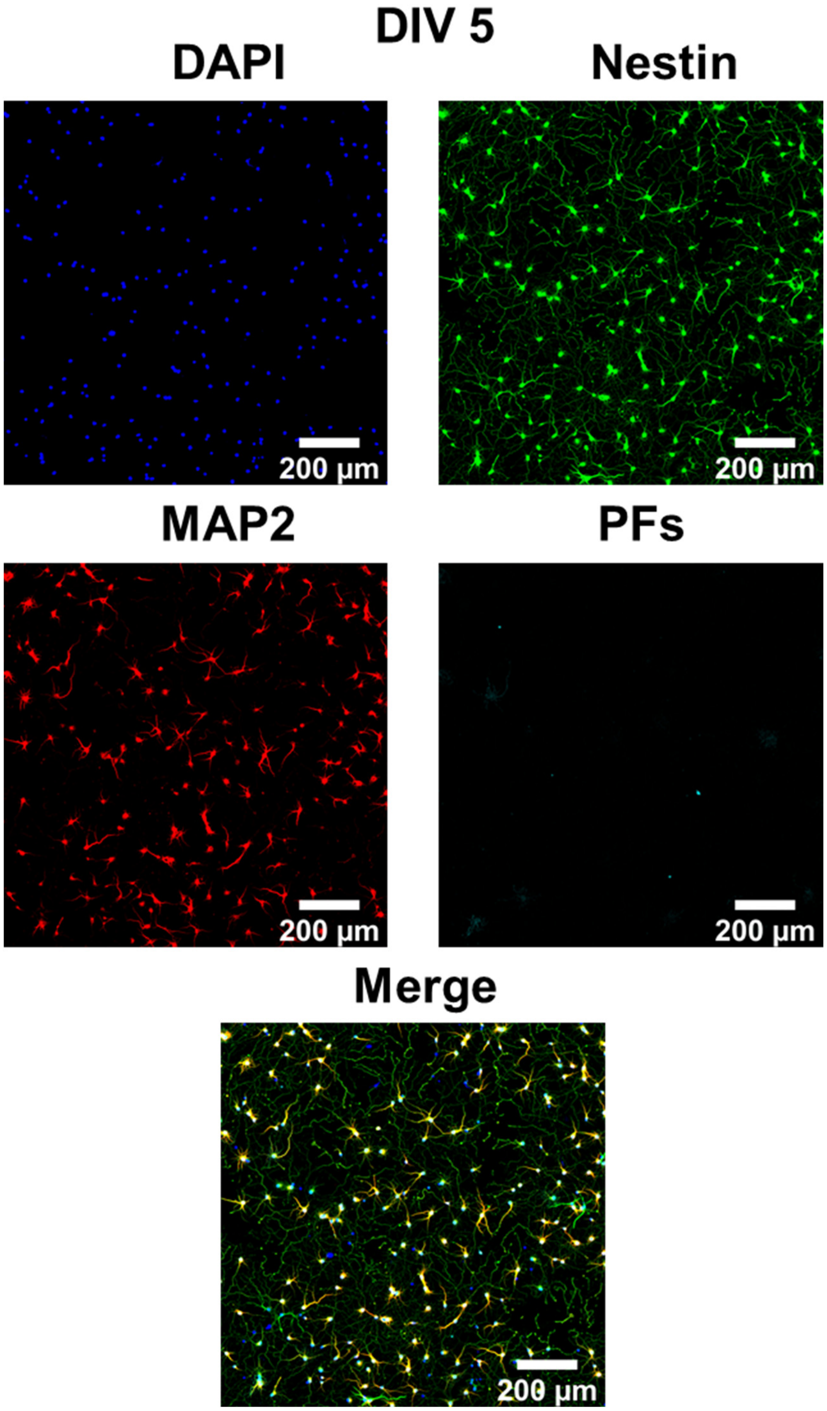


**Figure S10.** Individual channels of 3x3 tiled 20X magnification confocal fluorescence images (only merged image presented in the main text) of cultured hippocampal neurons after 1 hour incubation with negatively charged PFs (cyan) at DIV 5, co-stained with MAP2 (red, neuronal cell marker, specific to neuron cells), Nestin (green, progenitor cell marker, stains both neurons and glial cells) and DAPI (blue, nucleus staining).

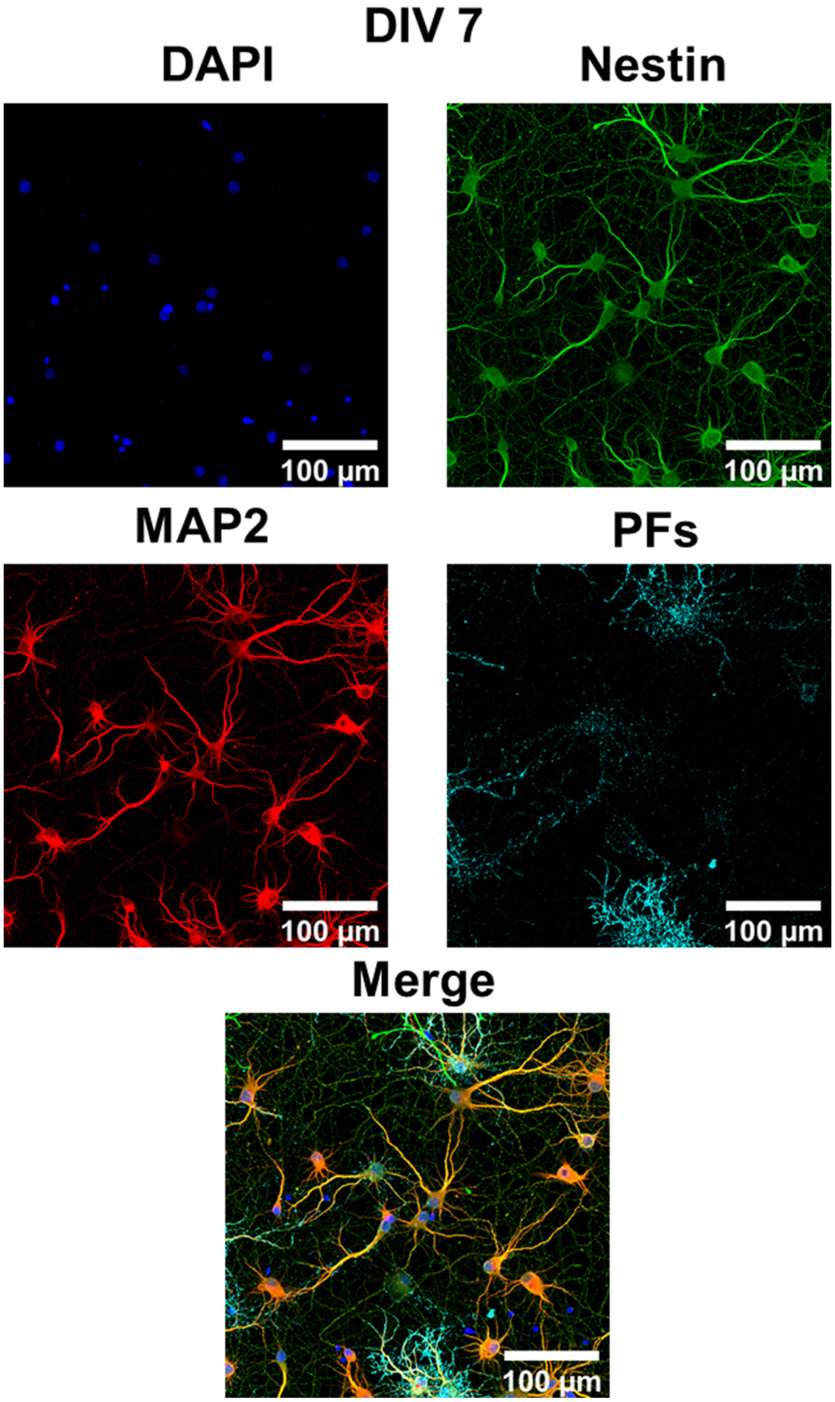


**Figure S11.** Individual channels of 20X magnification confocal fluorescence images (only merged image presented in the main text) of cultured hippocampal neurons after 1 hour incubation with negatively charged PFs (cyan) at DIV 7, co-stained with MAP2 (red, neuronal cell marker, specific to neuron cells), Nestin (green, progenitor cell marker, stains both neurons and glial cells) and DAPI (blue, nucleus staining).

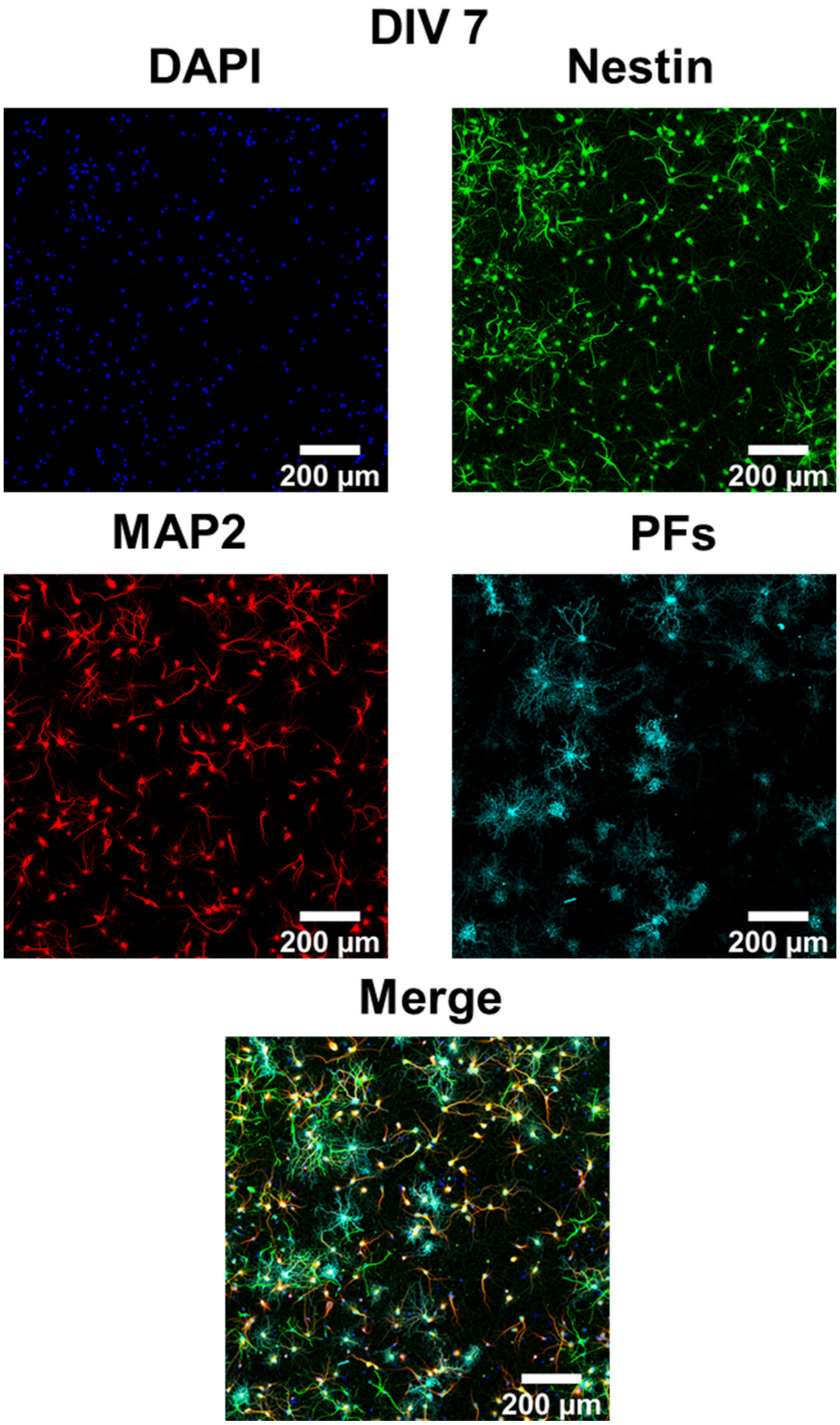


**Figure S12.** Individual channels of 3x3 tiled 20X magnification confocal fluorescence images (only merged image presented in the main text) of cultured hippocampal neurons after 1 hour incubation with negatively charged PFs (cyan) at DIV 7, co-stained with MAP2 (red, neuronal cell marker, specific to neuron cells), Nestin (green, progenitor cell marker, stains both neurons and glial cells) and DAPI (blue, nucleus staining).

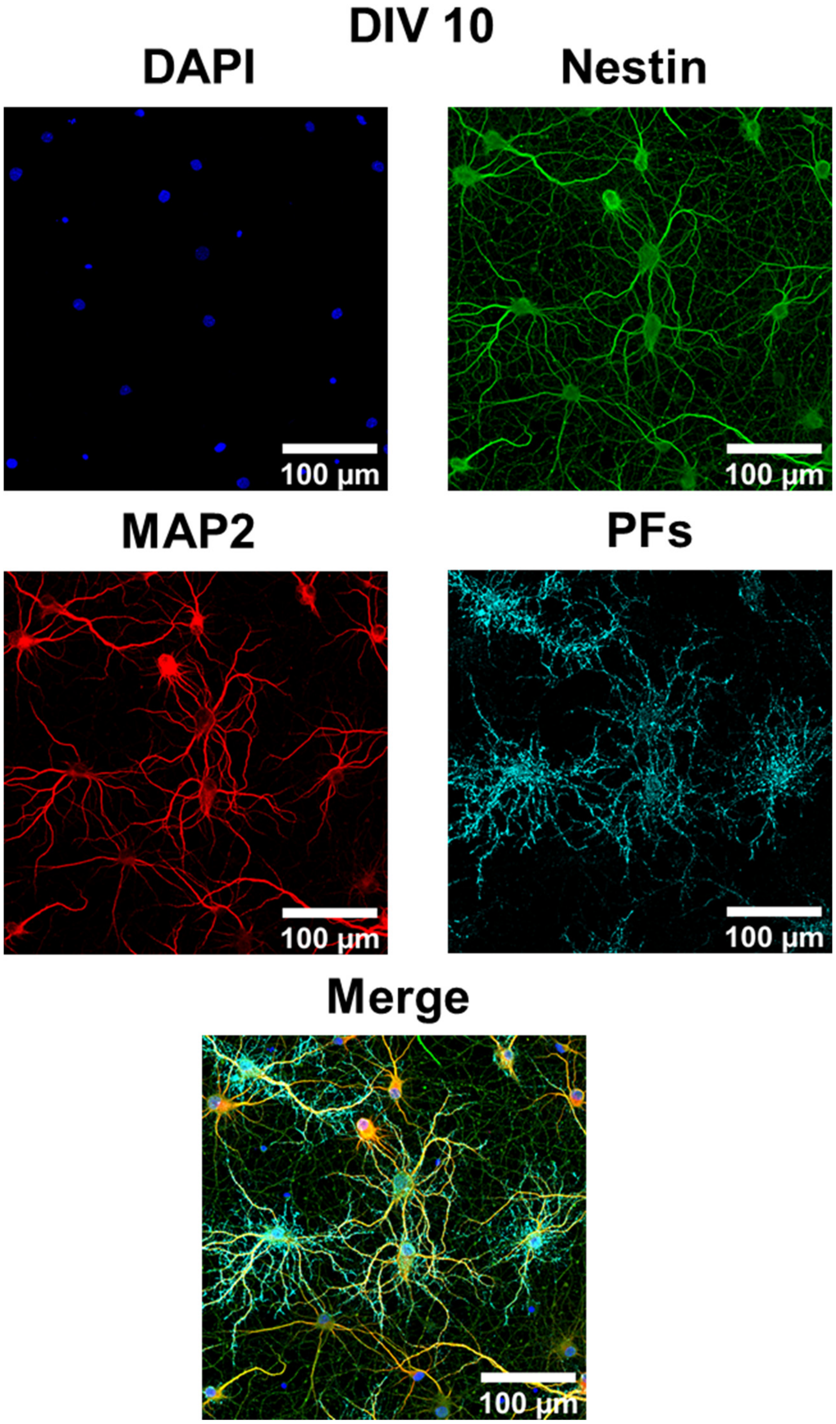


**Figure S13.** Individual channels of 20X magnification confocal fluorescence images (only merged image presented in the main text) of cultured hippocampal neurons after 1 hour incubation with negatively charged PFs (cyan) at DIV 10, co-stained with MAP2 (red, neuronal cell marker, specific to neuron cells), Nestin (green, progenitor cell marker, stains both neurons and glial cells) and DAPI (blue, nucleus staining).

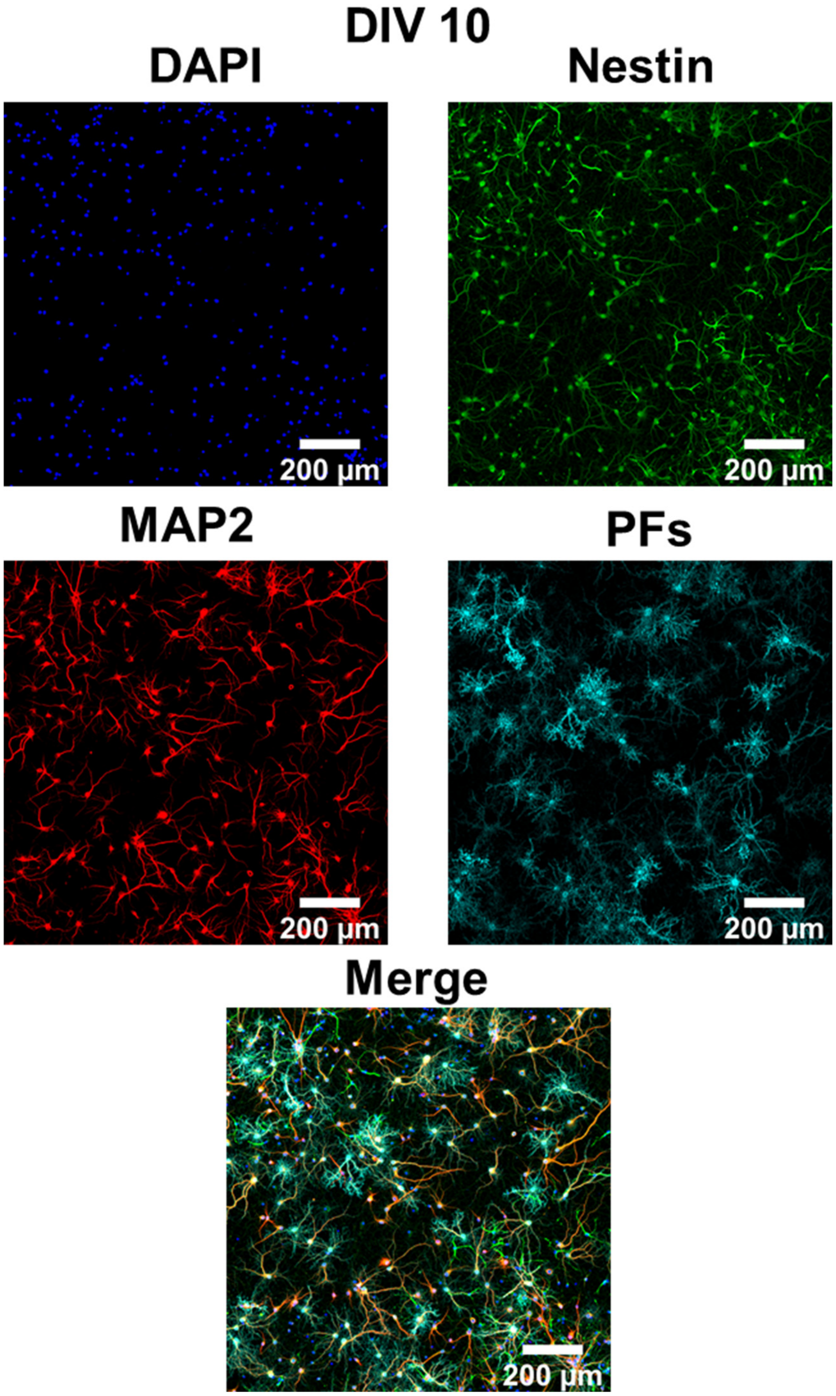


**Figure S14.** Individual channels of 3x3 tiled 20X magnification confocal fluorescence images (only merged image presented in the main text) of cultured hippocampal neurons after 1 hour incubation with negatively charged PFs (cyan) at DIV 10, co-stained with MAP2 (red, neuronal cell marker, specific to neuron cells), Nestin (green, progenitor cell marker, stains both neurons and glial cells) and DAPI (blue, nucleus staining).

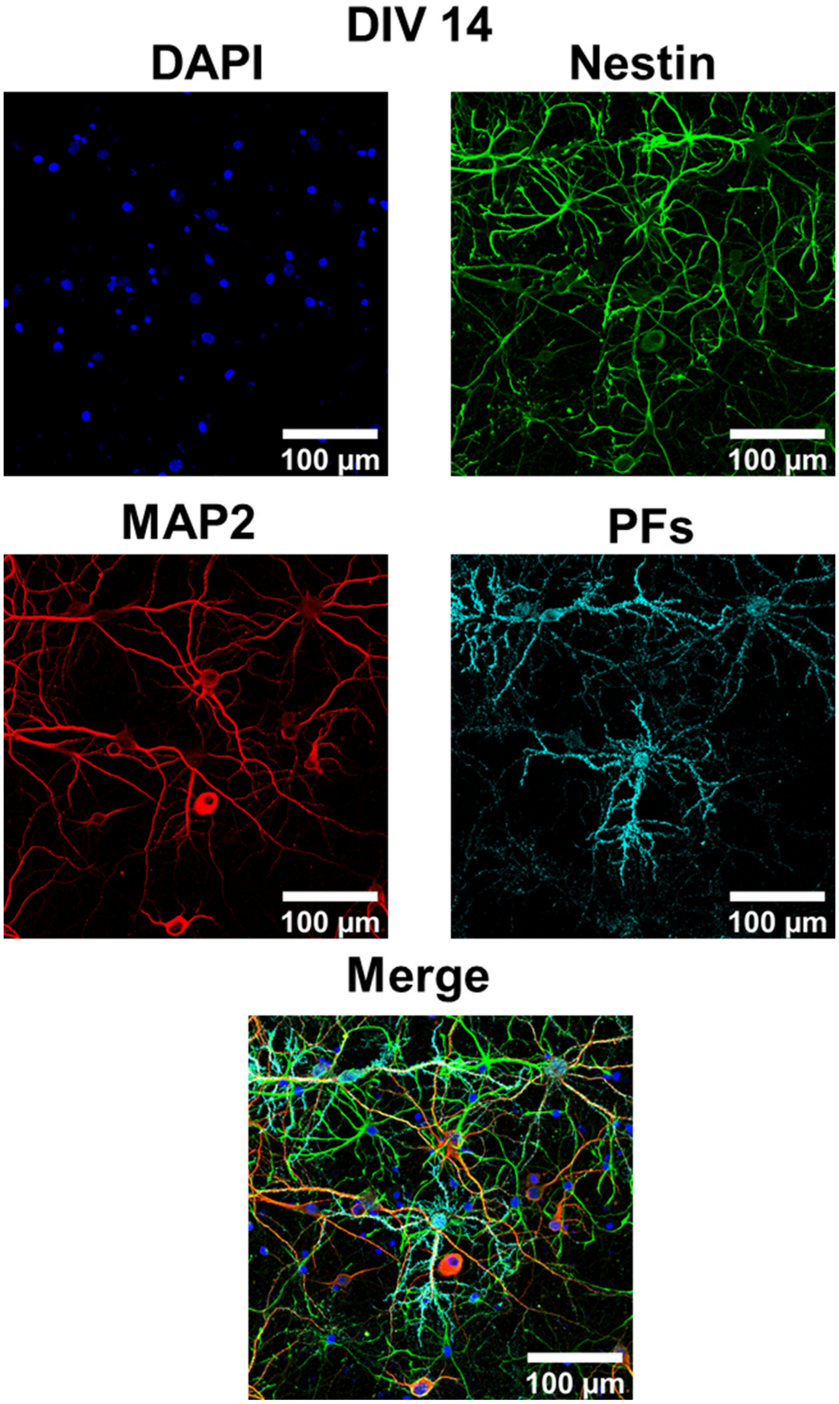


**Figure S15.** Individual channels of 20X magnification confocal fluorescence images (only merged image presented in the main text) of cultured hippocampal neurons after 1 hour incubation with negatively charged PFs (cyan) at DIV 14, co-stained with MAP2 (red, neuronal cell marker, specific to neuron cells), Nestin (green, progenitor cell marker, stains both neurons and glial cells) and DAPI (blue, nucleus staining).

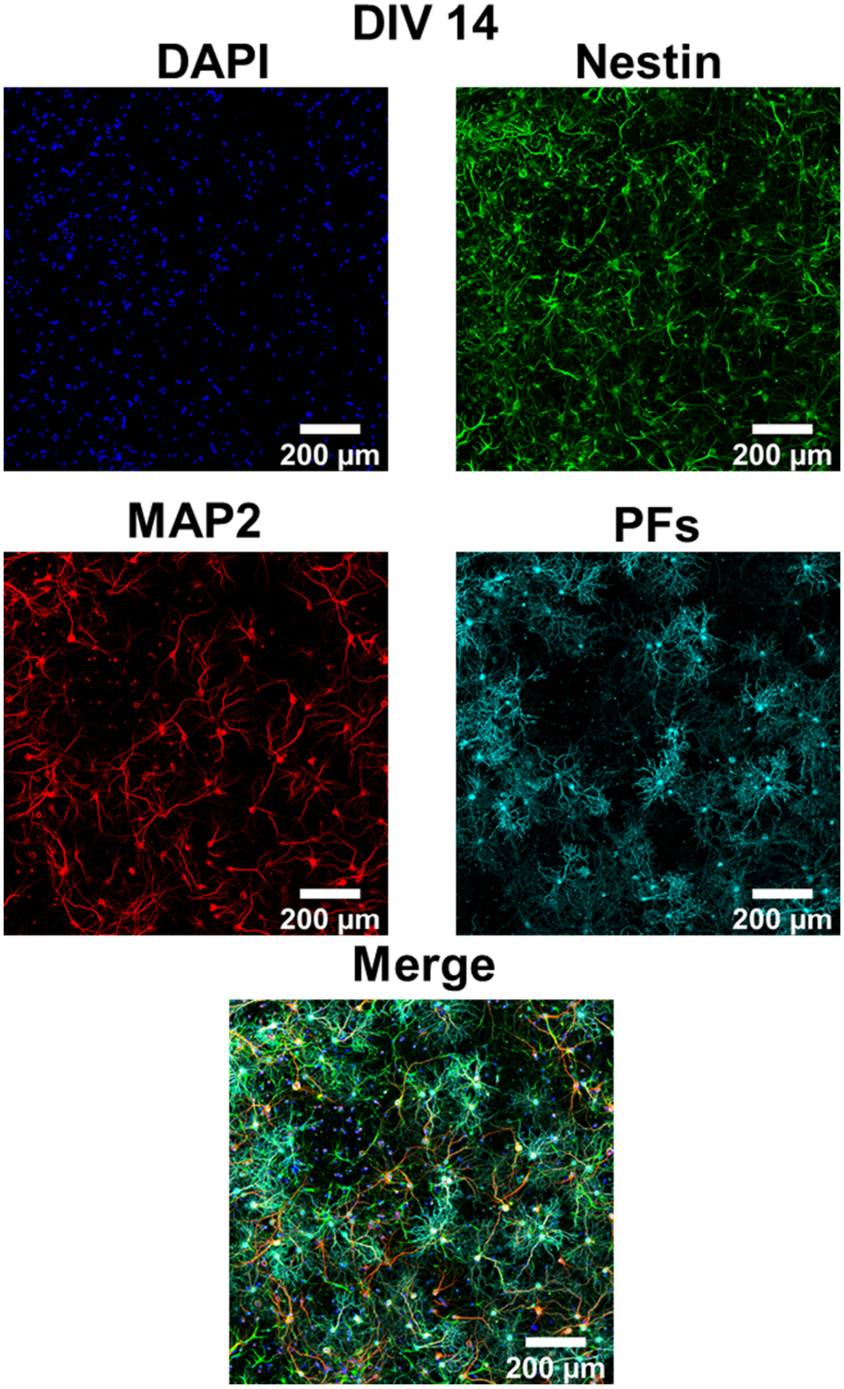


**Figure S16.** Individual channels of 3x3 tiled 20X magnification confocal fluorescence images (only merged image presented in the main text) of cultured hippocampal neurons after 1 hour incubation with negatively charged PFs (cyan) at DIV 14, co-stained with MAP2 (red, neuronal cell marker, specific to neuron cells), Nestin (green, progenitor cell marker, stains both neurons and glial cells) and DAPI (blue, nucleus staining).

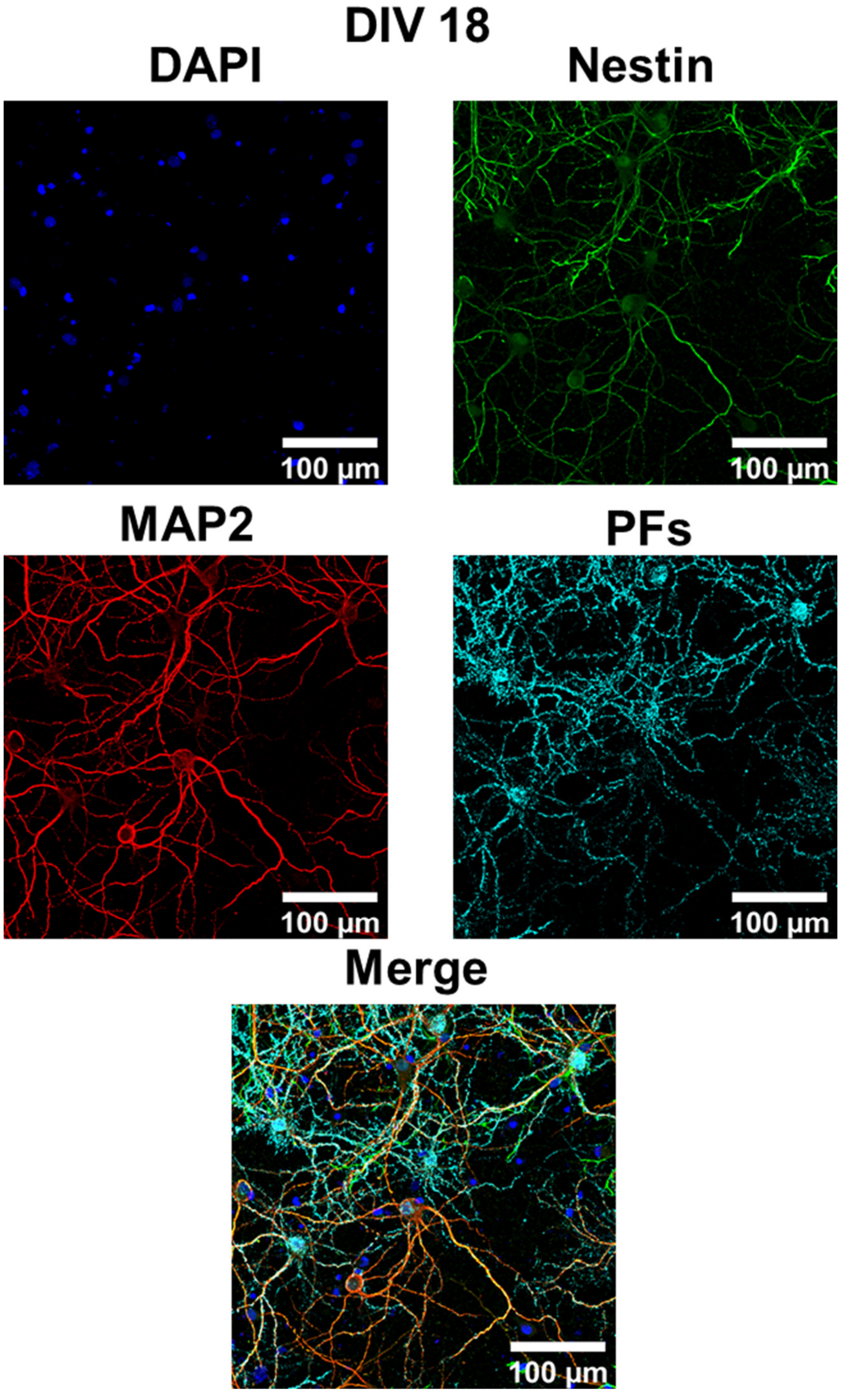


**Figure S17.** Individual channels of 20X magnification confocal fluorescence images (only merged image presented in the main text) of cultured hippocampal neurons after 1 hour incubation with negatively charged PFs (cyan) at DIV 18, co-stained with MAP2 (red, neuronal cell marker, specific to neuron cells), Nestin (green, progenitor cell marker, stains both neurons and glial cells) and DAPI (blue, nucleus staining).

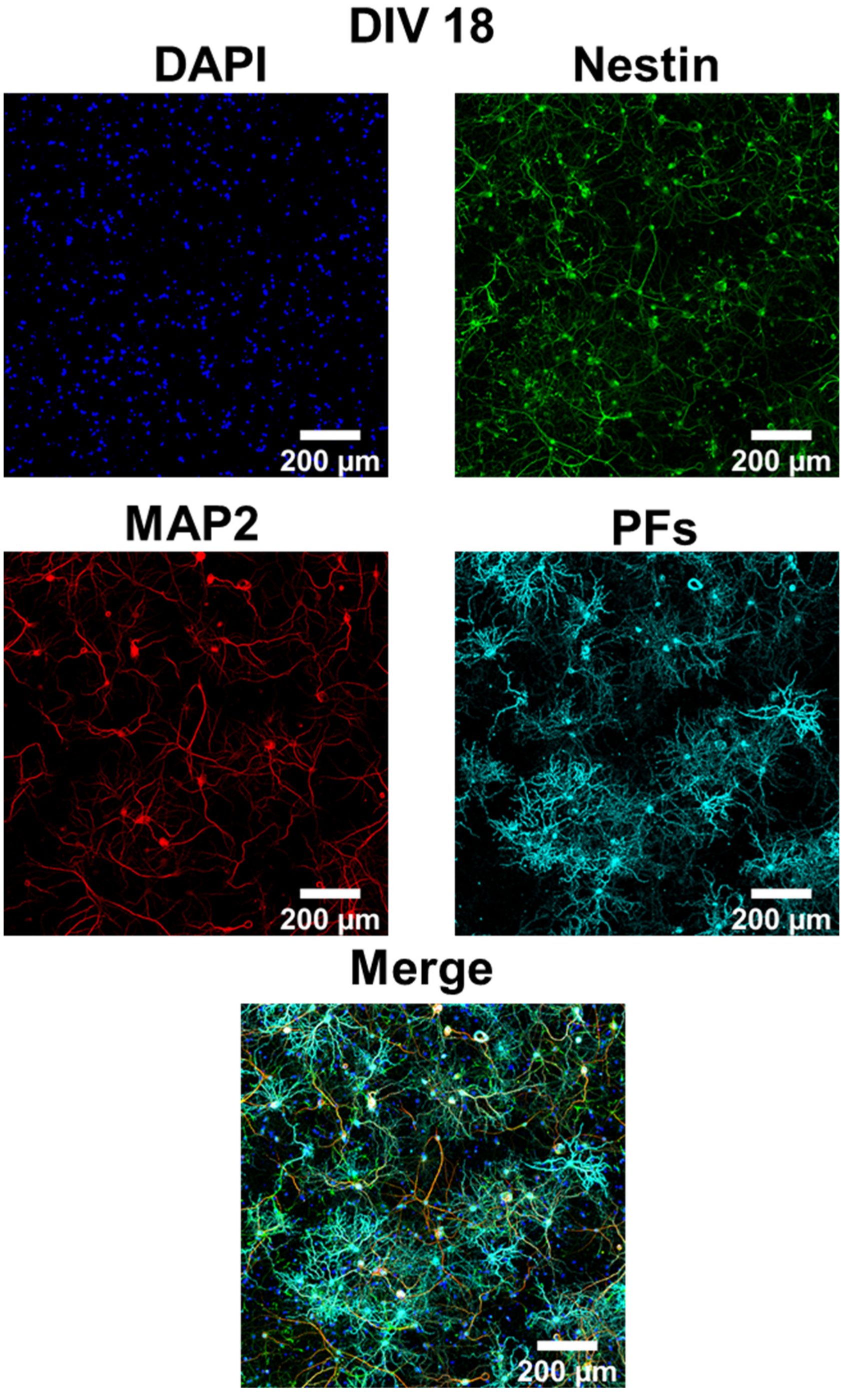


**Figure S18.** Individual channels of 3x3 tiled 20X magnification confocal fluorescence images (only merged image presented in the main text) of cultured hippocampal neurons after 1 hour incubation with negatively charged PFs (cyan) at DIV 18, co-stained with MAP2 (red, neuronal cell marker, specific to neuron cells), Nestin (green, progenitor cell marker, stains both neurons and glial cells) and DAPI (blue, nucleus staining).

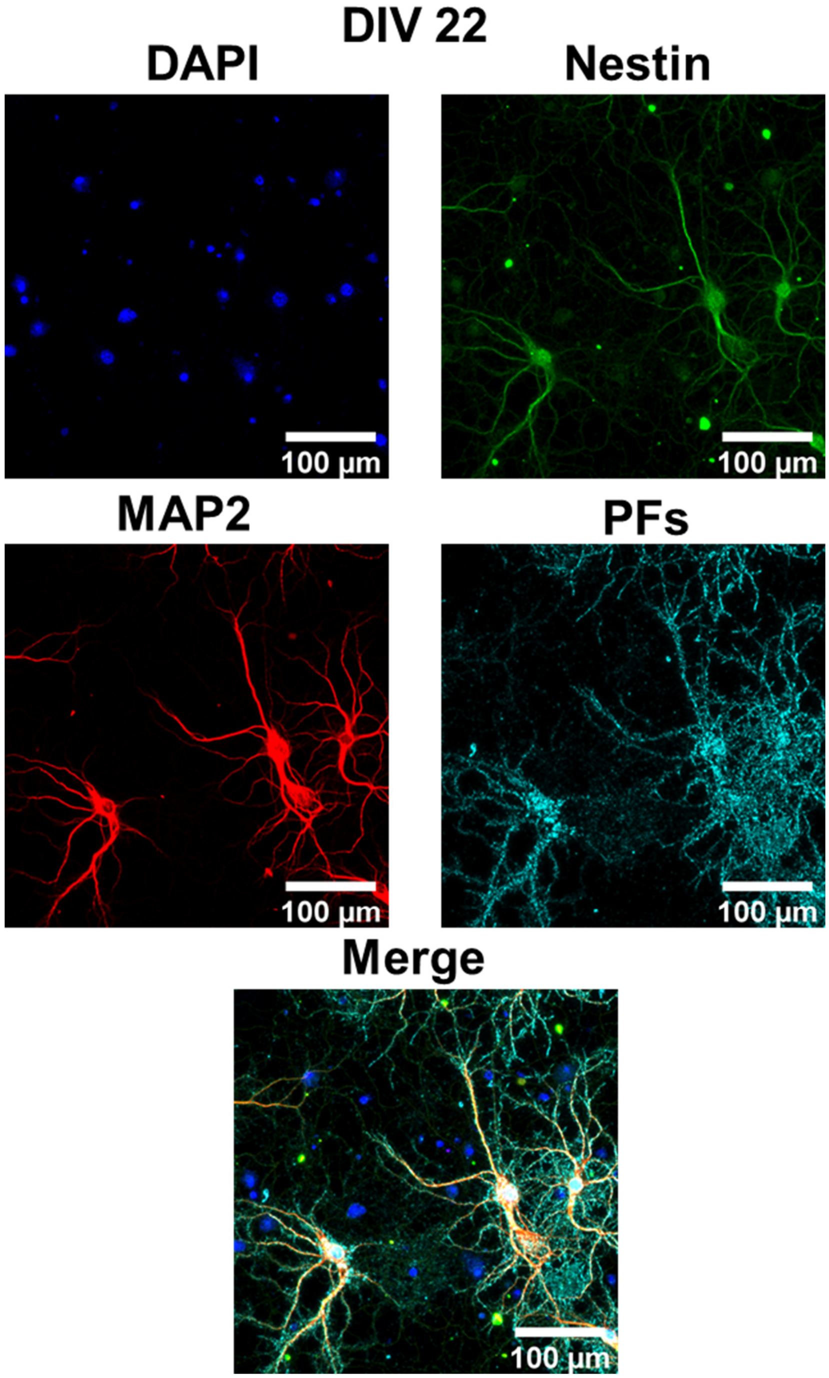


**Figure S19.** Individual channels of 20X magnification confocal fluorescence images (only merged image presented in the main text) of cultured hippocampal neurons after 1 hour incubation with negatively charged PFs (cyan) at DIV 22, co-stained with MAP2 (red, neuronal cell marker, specific to neuron cells), Nestin (green, progenitor cell marker, stains both neurons and glial cells) and DAPI (blue, nucleus staining).

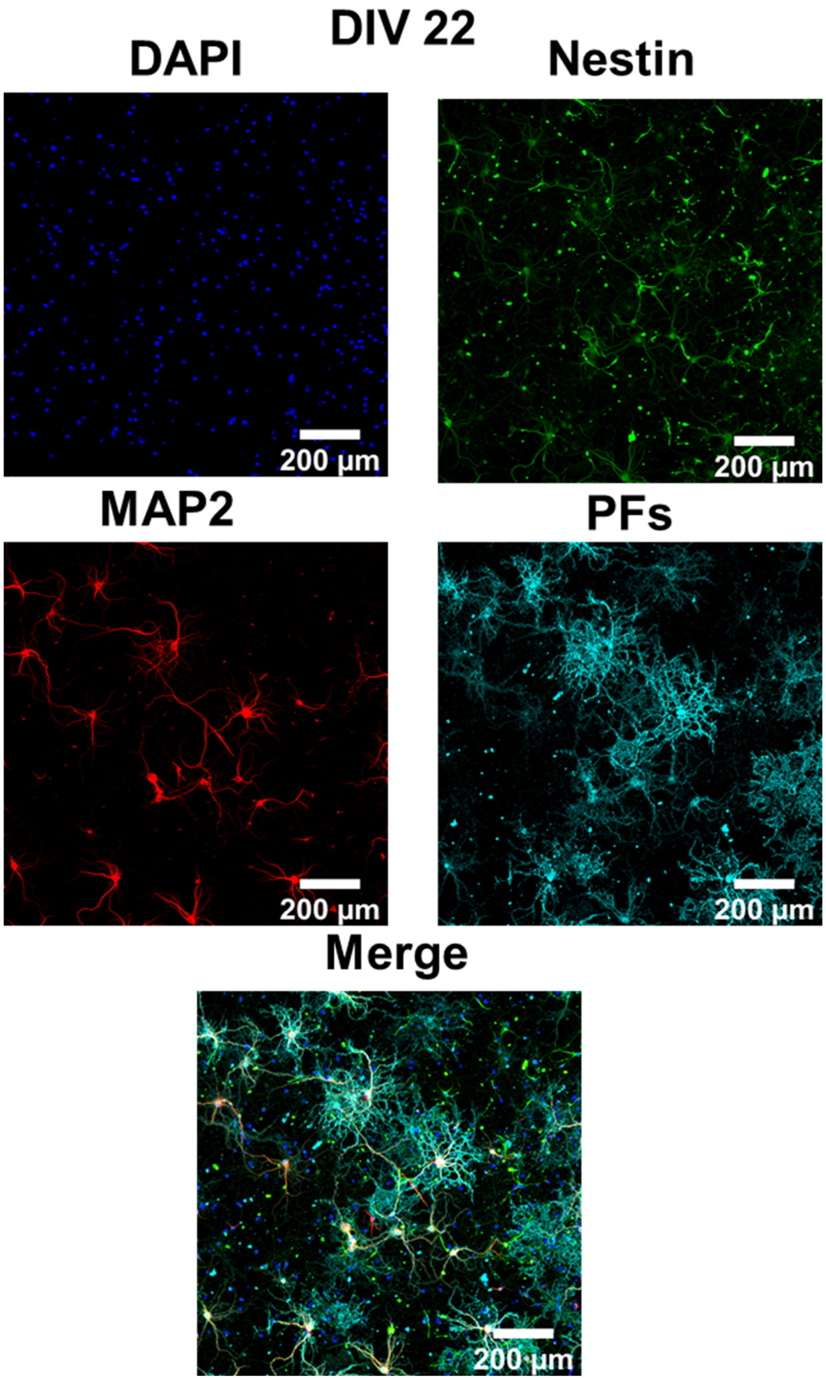


**Figure S20.** Individual channels of 3x3 tiled 20X magnification confocal fluorescence images (only merged image presented in the main text) of cultured hippocampal neurons after 1 hour incubation with negatively charged PFs (cyan) at DIV 22, co-stained with MAP2 (red, neuronal cell marker, specific to neuron cells), Nestin (green, progenitor cell marker, stains both neurons and glial cells) and DAPI (blue, nucleus staining)

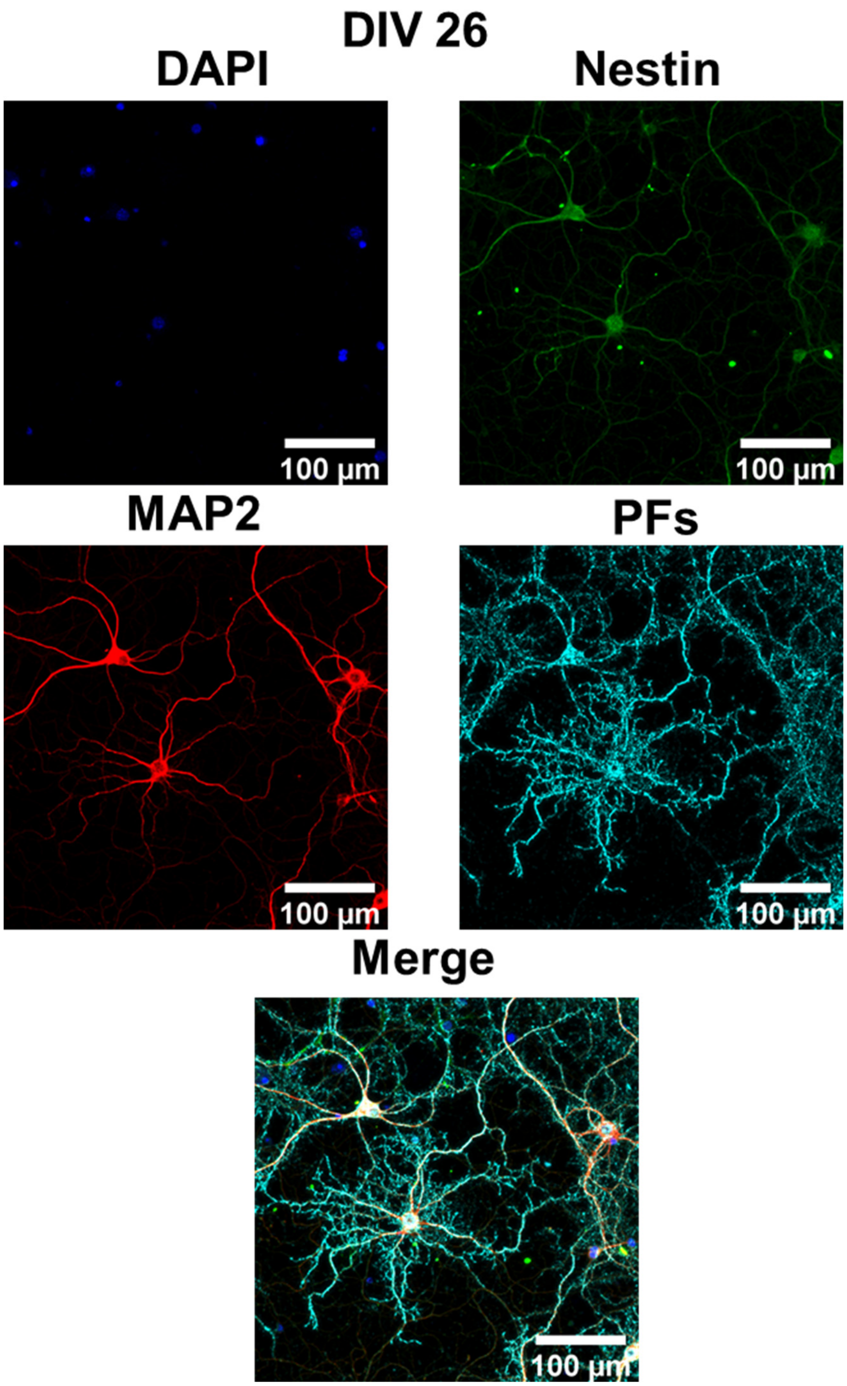


**Figure S21.** Individual channels of 20X magnification confocal fluorescence images (only merged image presented in the main text) of cultured hippocampal neurons after 1 hour incubation with negatively charged PFs (cyan) at DIV 26, co-stained with MAP2 (red, neuronal cell marker, specific to neuron cells), Nestin (green, progenitor cell marker, stains both neurons and glial cells) and DAPI (blue, nucleus staining).

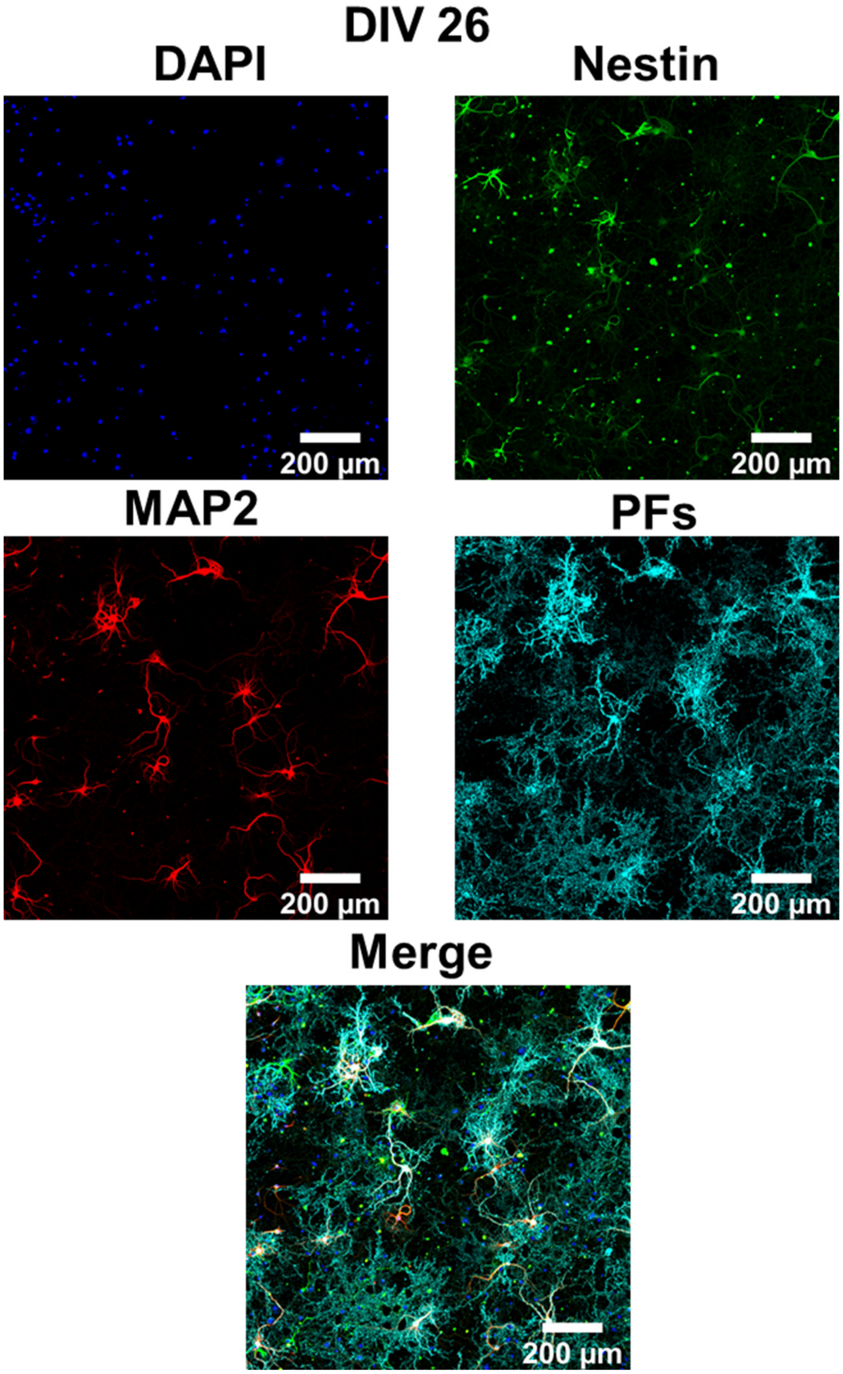


**Figure S22.** Individual channels of 3x3 tiled 20X magnification confocal fluorescence images (only merged image presented in the main text) of cultured hippocampal neurons after 1 hour incubation with negatively charged PFs (cyan) at DIV 26, co-stained with MAP2 (red, neuronal cell marker, specific to neuron cells), Nestin (green, progenitor cell marker, stains both neurons and glial cells) and DAPI (blue, nucleus staining).

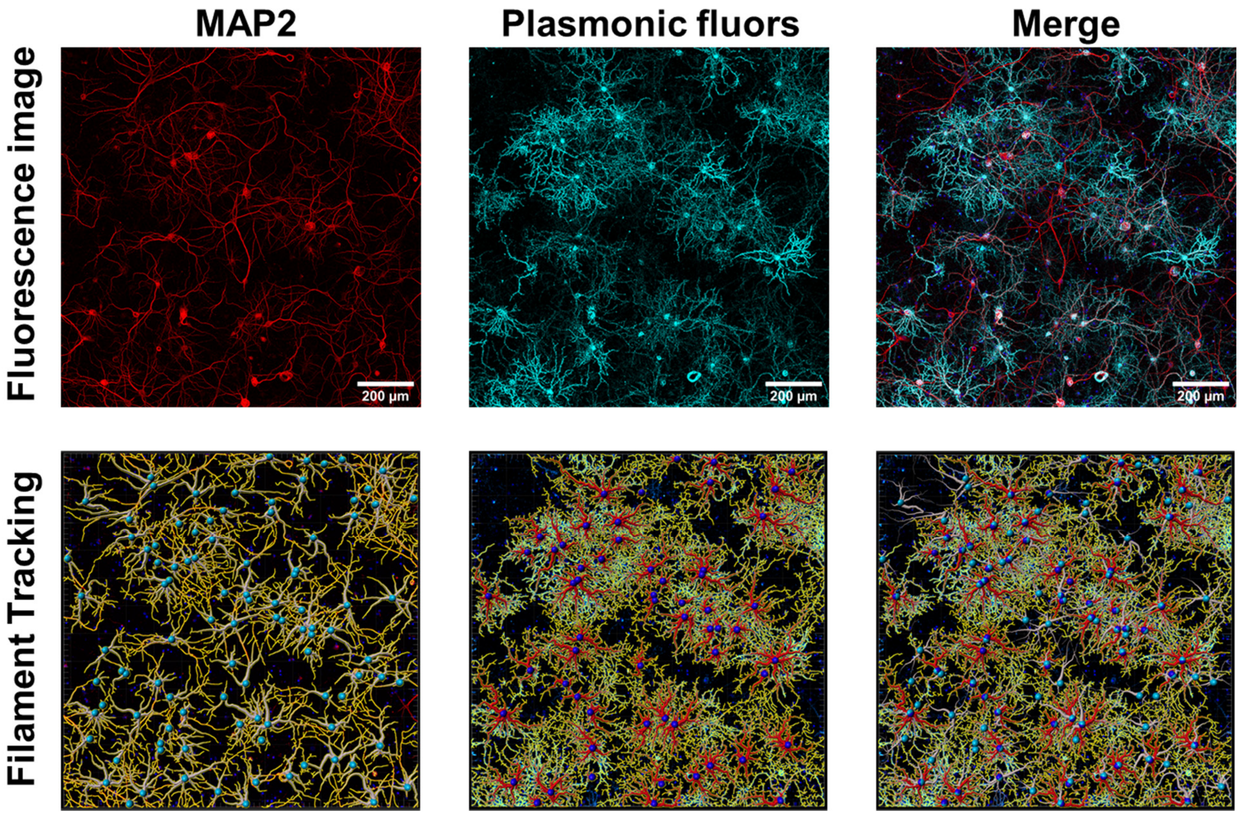


**Figure S23.** Top panel exhibits individual channels of 20X magnification confocal fluorescence image of cultured hippocampal neurons after 1 hour incubation with negatively charged PFs (cyan) at DIV 18, co-stained with MAP2 (red, neuronal cell marker, specific to neuron cells), Nestin (green, progenitor cell marker, stains both neurons and glial cells) and DAPI (blue, nucleus staining). Bottom panel represents corresponding filament traced images generated via filament tracer module of IMARIS software (OXFORD INSTRUMENTS) for MAP2 and PFs channel.

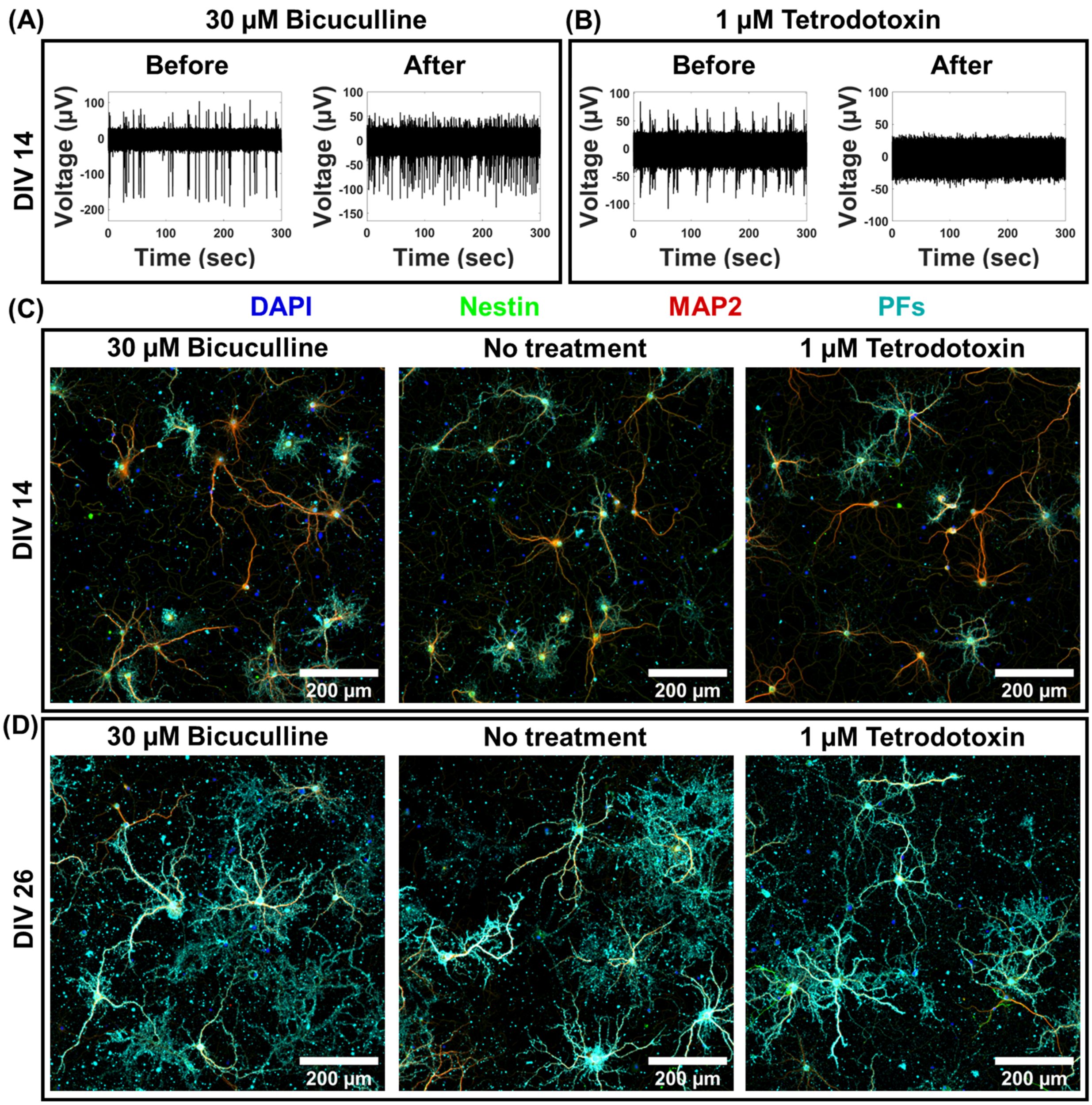


**Figure S24.** A single trace of spike recording before and after neurons were incubated with (A) 30μM bicuculline and (B) 1 μM tetrodotoxin for 15 minutes. The traces are representative ones from a total of 32 active channels measured from primary cultured hippocampal neurons cultured on a microelectrode array (MEA). The experiment was repeated tw0 times independently with similar results. Confocal fluorescence images of cultured hippocampal neurons after pharmacologically manipulating the electrophysiological activity and subsequent 1 hour incubation with negatively charged PFs (cyan) at (C) DIV 14 and (D) DIV 26, co-stained with MAP2 (red, neuronal cell marker, specific to neuron cells), Nestin (green, progenitor cell marker, stains both neurons and glial cells) and DAPI (blue, nucleus staining). Each block shows the 3×3 tiled image obtained from 20X magnification images. (n=2 independent experiments).
